# Cannabidiol (CBD) repurposing as antibacterial: promising therapy of CBD plus polymyxin B against superbugs

**DOI:** 10.1101/2021.04.12.439341

**Authors:** Nathália Abichabki, Luísa V. Zacharias, Natália C. Moreira, Fernando Bellissimo-Rodrigues, Fernanda L. Moreira, Jhohann R. L. Benzi, Tânia M. C. Ogasawara, Joseane C. Ferreira, Leonardo R. L. Pereira, Gilberto Ú. L. Braga, Camila M. Ribeiro, Fernando R. Pavan, Antonio W. Zuardi, Jaime E. C. Hallak, José A. S. Crippa, Vera L. Lanchote, Rafael Cantón, Ana Lúcia C. Darini, Leonardo N. Andrade

## Abstract

Abstract

Multidrug-resistant (MDR) and extensively drug-resistant (XDR) bacteria are a major worldwide public health problem. In the last decades, resistance to last-resort antibiotics such as polymyxin B (PB) have been increasingly observed among these superbugs, compromising the effectiveness of antimicrobial therapy. The present study aimed (i) to assess the ultrapure Cannabidiol (CBD) antibacterial activity against a broad diversity of Gram-negative (GN) and Gram-positive (GP) bacteria (44 different species, 95 strains), comprising standard strains and clinical isolates, and (ii) to investigate the antibacterial activity of the combination CBD + PB against GN bacteria, including chromosomal- and plasmid-acquired PB-resistant and intrinsically PB-resistant GNB. We evaluated CBD *in vitro* antibacterial activity using the standard broth microdilution method, and the antibacterial activity of the combination CBD + PB was screened using the standard broth microdilution and confirmed by checkerboard assay. CBD exhibited antibacterial activity against different GP bacterial species, lipooligosaccharide (LOS)-expressing GN diplococcus (GND) (*Neisseria gonorrhoeae*, *Neisseria meningitidis*, and *Moraxella catarrhalis*), and *Mycobacterium tuberculosis*. The combination CBD + PB exhibited antibacterial activity against PB-resistant GNB (e.g., *Klebsiella pneumoniae*) as well as additive and/or synergistic effect against LOS-expressing GND. The antibacterial activity of the combination CBD + PB against *Pseudomonas aeruginosa* and plasmid-mediated colistin-resistant (MCR-1) *E. coli* strains could be only demonstrated in the presence of phenylalanine-arginine-β-naphthylamide (PAβN). In conclusion, our results show promising translational potential of the combination CBD + PB against MDR and XDR GNB, including PB-resistant *K. pneumoniae*, highlighting its potential as a rescue treatment for life-threatening infections caused by these superbugs.

**One Sentence Summary:** Promising combination of cannabidiol (CBD) + polymyxin B (PB) against superbugs (e.g., PB-resistant Gram-negative bacilli): Repurposing CBD

## INTRODUCTION

Antimicrobial resistance (AMR) is a major worldwide public health problem, mainly due to the increasing incidence of bacterial infections caused by multidrug-resistant (MDR) and extensively drug-resistant (XDR) Gram-negative bacilli (GNB). A group of pathogens identified by the acronym ESKAPE (*Enterococcus faecium*, *Staphylococcus aureus*, *Klebsiella pneumoniae*, *Acinetobacter baumannii*, *Pseudomonas aeruginosa*, and *Enterobacter* spp.) either resistant to third-generation cephalosporin and/or carbapenem, and to vancomycin and/or methicillin is of special a concern (*1*).

In this context, polymyxin B (PB) was reintroduced into the clinical practice and recognized as reserve group antibiotics (last-resort) to treat serious infections due to third-generation cephalosporin- or carbapenem-resistant GNB. However, acquired resistance to polymyxins (including chromosomal- and plasmid-mediated resistance) has been increasingly detected in several GNB such as *Enterobacterales* species (e.g., *K. pneumoniae*) and non-fermenting (NF) GNB (e.g., *A. baumannii* and *P. aeruginosa*) (*2, 3*).

The antibacterial activity of polymyxins is due to an electrostatic interaction between the positively charged polymyxin and the phosphate groups of the negatively charged lipid A, on lipopolysaccharide (LPS) or lipooligosaccharide (LOS), in the outer membrane of Gram-negative (GN) bacteria. Thus, the LPS or LOS is destabilized leading to bacterial cell envelop disruption. Polymyxin-resistant GN bacteria usually have the addition of phosphoethanolamine (pEtN) and/or 4-amino-4-deoxy-L-arabinose (L-Ara4N) cationic groups on their lipid A molecule, giving it positive charges and resulting in the repulsion of polymyxin molecule (*2, 4*).

Considering health, social and economic implications of growing AMR, the World Health Organization (WHO) reported a global priority list of antibiotic-resistant bacteria, including ESKAPE pathogens, to guide research, discovery, and development (R&D) of new antibiotics (*5, 6*).

In this context, many substances have been investigated regarding their antimicrobial activity, including natural products. Cannabidiol (CBD) is the major non-psychoactive component isolated from *Cannabis sativa* and has been associated with multiple and potential biological activities, especially anxiolytic, antipsychotic, anti-inflammatory, analgesic, antioxidant and neuroprotective properties (*7–10*). Some studies also describe the potential of CBD to inhibit the spread of cancerous cells (*11, 12*). Since the 1950’s, *C. sativa*-based preparations have been investigated regarding their antibacterial activity (*13–15*). CBD was also described as an inhibitor of membrane vesicles released from GN bacteria (*16*). Despite this fact, few studies described the antibacterial activity of ultrapure CBD against Gram-positive (GP) bacteria and its property of inhibiting biofilms formation and eradicate preformed biofilms (*10, 13, 14, 17–19*). In addition, few GNB species or genus as well as different phenotype/genotype of resistance have been evaluated, and the combination with antibiotics such as PB was poorly explored, particularly against MDR and XDR GNB, commonly called superbugs (*20, 21*).

The present study aimed to evaluate the *in vitro* (i) antibacterial activity of ultrapure CBD against a broad diversity of bacteria (44 different species, 95 strains), comprising standard strains and clinical isolates (including MDR ESKAPE pathogens), and (ii) antibacterial activity of the combination CBD + PB against GN bacteria, including chromosomal- and plasmid-acquired PB- resistant and intrinsically PB-resistant GNB (11 species, 51 strains).

## RESULTS

### CBD antibacterial activity against GP and GN bacteria, and *Mycobacterium tuberculosis*

CBD showed different level of antibacterial activity (minimal inhibitory concentration [MIC]) against all 13 different species of GP bacteria (21 strains) evaluated, including susceptible and MDR strains: MIC = 2 µg/mL for *E. faecium* (n=2); MIC = 4 µg/mL for *Enterococcus* spp. (n=4), *Staphylococcus* spp. (n=10), *Micrococcus luteus* (n=1), and *Rhodococcus equi* (n=1); MIC = 32 µg/mL for *Streptococcus pyogenes* (n=1) and *Streptococcus pneumoni*ae (n=1); and MIC = 64 µg/mL *Streptococcus agalactiae* (n=1) (Table S1).

The antibacterial activity of CBD was also observed against LOS-expressing GN diplococcus (GND), *Moraxella catarrhalis* ATCC 25238 (MIC = 64 µg/mL), *Neisseria meningitidis* ATCC 13077 (MIC = 128 µg/mL), and *Neisseria gonorrhoeae* ATCC 19424 (MIC = 256 µg/mL); as well as against *Mycobacterium tuberculosis* H37Rv (MIC = 9,37±1,88 µg/mL) and MDR *M. tuberculosis* CF86 (MIC = 18,78±5,95 µg/mL) (Table S1).

We observed no difference between MIC and minimum bactericidal concentration (MBC) values among susceptible and MDR strains evaluated.

For *S. aureus* ATCC 29213, we observed higher CBD MIC (64 µg/mL) when the assay was performed using MH-F broth (5% lysed horse blood + 0.1% β-Nicotinamide adenine dinucleotide [β-NAD] 20 mg/mL), in comparison with standard protocols using CAMHB for *S. aureus* (CBD MIC = 4 µg/mL) (Fig. S1) (*22, 23*).

CBD MIC values for all GNB evaluated were higher than 256 µg/mL (Table S1). For *E. coli* ATCC 25922, *K. pneumoniae* ATCC 13883, *A. baumannii* ATCC 19606, and *P. aeruginosa* ATCC 27853 higher CBD concentrations were also evaluated, and again no antibacterial activity was observed up to 8,192 µg/mL.

Complementarily, we did not detect any antibacterial activity of CBD against GNB in the presence of efflux pump inhibitors (phenylalanine-arginine-β-naphthylamide [PAβN], reserpine, or curcumin).

### Antibacterial activity of CBD in combination with PB (CBD + PB) against GN bacteria

#### Broth Microdilution method with fixed concentration of CDB (256 µg/mL)

The antibacterial activity of the combination CBD + PB was observed against 11 different species (51 strains) of GN bacteria, including standard strains and clinical isolates (Table 1).

**Table 1.** PB concentrations in the combination of CBD (256 µg/mL) + PB minimal antibiotic concentration (MAC) compared to PB MIC. Compared to PB MIC, there is a reduction in the PB concentration required to the antibacterial activity of CBD + PB. These PB concentrations represent the PB MAC and allow CBD antibacterial activity.

| Strain | PB (µg/mL) |  |
| --- | --- | --- |
|  | MAC |  |
|  | in the combination | MIC |
|  | CBD + PB |  |
| *K. pneumoniae* ATCC 13883<sup>T</sup> | 0.125 | 8 |
| *K. pneumoniae* ATCC BAA-1705 | 0.25 | 0.5 |
| *K. pneumoniae* NCTC 13443 | 0.06 | 1 |
| *K. pneumoniae* C9<sup>§</sup> | ≤ 1 | 64 |
| *K. pneumoniae* D1<sup>§</sup> | 0.25 | 1 |
| *K. pneumoniae* RP 62<sup>‡</sup> | ≤ 0.5 | 32 |
| *K. pneumoniae* L1<sup>γ</sup> | ≤ 0.25 | 8 |
| *K. pneumoniae* L2<sup>γ</sup> | ≤ 0.5 | 64 |
| *K. pneumoniae* L3<sup>γ</sup> | ≤ 0.25 | 64 |
| *K. pneumoniae* L5<sup>γ</sup> | ≤ 0.03 | 1 |
| *K. pneumoniae* L8<sup>γ</sup> | ≤ 0.25 | 256 |
| *K. pneumoniae* L9<sup>γ</sup> | ≤ 0.25 | 32 |
| *K. pneumoniae* L12<sup>γ</sup> | 0.06 | 2 |
| *K. pneumoniae* L13<sup>y</sup> | $\leq 0.25$ | 16 |
| *K. pneumoniae* L14<sup>y</sup> | $\leq 0.25$ | 32 |
| *K. pneumoniae* L15<sup>y</sup> | 0.5 | 4 |
| *K. pneumoniae* L16<sup>y</sup> | 0.5 | 4 |
| *K. pneumoniae* L17<sup>y</sup> | $\leq 0.5$ | 64 |
| *K. pneumoniae* L18<sup>y</sup> | 8 | 64 |
| *K. pneumoniae* L19<sup>y</sup> | 1 | 16 |
| *K. pneumoniae* L22<sup>y</sup> | $\leq 0.5$ | 256 |
| *K. pneumoniae* L26<sup>y</sup> | $\leq 0.5$ | 64 |
| *K. pneumoniae* L27<sup>y</sup> | 0.06 | 2 |
| *K. pneumoniae* L28<sup>y</sup> | $\leq 0.5$ | 64 |
| *K. pneumoniae* L29<sup>y</sup> | $\leq 0.25$ | 16 |
| *K. pneumoniae* L30<sup>y</sup> | 0.25 | 1 |
| *K. pneumoniae* L31<sup>y</sup> | 0.06 | 8 |
| *K. pneumoniae* L33<sup>y</sup> | 0.06 | 16 |
| *K. pneumoniae* L34<sup>y</sup> | $\leq 0.125$ | 32 |
| *K. quasipneumoniae* ATCC 700603 | 0.125 | 0.5 |
| *E. cloacae* ATCC 13047<sup>T</sup> | $\leq 0.5$ | 256 |
| *E. coli* ATCC 25922 | 0.06 | 0.5 |
| *E. coli* CTX-M-15 | 0.06 | 0.5 |
| *E. coli* RP62T<sup>†</sup> | $\leq 0.01$ | 2 |
| *E. coli* 72H<sup>#</sup> | 1 | 2 |
| *E. coli* NCTC 13846 | 1 | 2 |
| *A. baumannii* ATCC 19606<sup>T</sup> | 0.25 | 1 |
| *A. baumannii* 136 SP<sup>*</sup> | 0.25 | 1 |
| *A. baumannii* L7<sup>γ</sup> | ≤ 0.03 | 2 |
| *A. baumannii* L21<sup>γ</sup> | 0.125 | 1 |
| *A. baumannii* L25<sup>γ</sup> | 0.125 | 1 |
| *A. baumannii* L35<sup>γ</sup> | 0.125 | 0.5 |
| *P. aeruginosa* ATCC 27853 | 1 | 1 |
| *P. aeruginosa* HC 103<sup>Φ</sup> | 1 | 2 |
| *P. aeruginosa* PAO1 | 0.5 | 1 |
| *P. aeruginosa* L36<sup>γ</sup> | 1 | 2 |
| *S. maltophilia* ATCC 13673<sup>T</sup> | 0.25 | 2 |
| *E. tarda* ATCC 15947<sup>T</sup> | 0.25 | > 8 |
| *M. catarrhalis* ATCC 25238 | ≤ 0.01 | 0.5 |
| *N. meningitidis* ATCC 13077 | ≤ 0.01 | 64 |
| *N. gonorrhoeae* ATCC 19424 | ≤ 0.03 | 16 |

CBD: Cannabidiol; PB: Polymyxin B; ATCC: *American Type Culture Collection*; <sup>T</sup>*type strain*; <sup>δ</sup>Palmeiro *et al*, 2019 (64); <sup>‡</sup>Andrade *et al*, 2014 (65); <sup>†</sup>Azide-resistant *E. coli*, transconjugant (IncFIIk-Tn4401-bla KPC-2) KPC producer, Andrade *et al*, 2014 (65).<sup>#</sup> Fernandes *et al*, 2016 (66); <sup>\*</sup>Clímaco *et al*, 2013 (67); <sup>Φ</sup>Galetti *et al*, 2015 (68); <sup>γ</sup>Clinical isolates, non MCR-1.

In the combination CBD (256 µg/mL) + PB (0.01 - 512 µg/mL), compared to PB MIC, we observed a minimal 3-fold reduction in the PB concentration required to exhibit antibacterial activity of CBD + PB against PB-susceptible GNB (NFGNB and *Enterobacterales*). For PB- susceptible strains, the combination CBD + PB against *K. pneumoniae* led to a 2-fold reduction in the PB concentration compared to PB MIC, while it was observed just 1-fold reduction for *P. aeruginosa* (Table 1 and Fig. 1F).

**Fig. 1.**
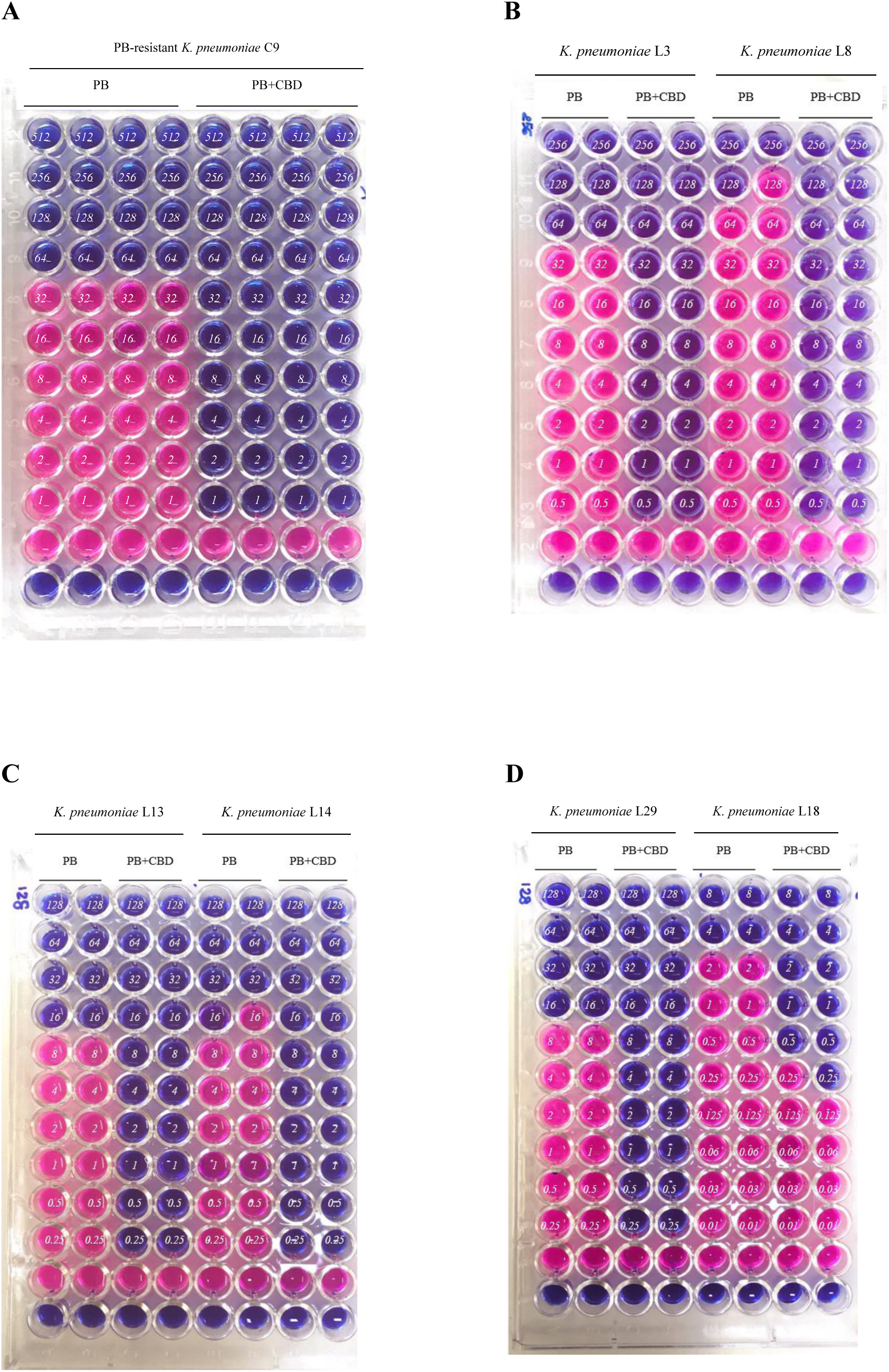
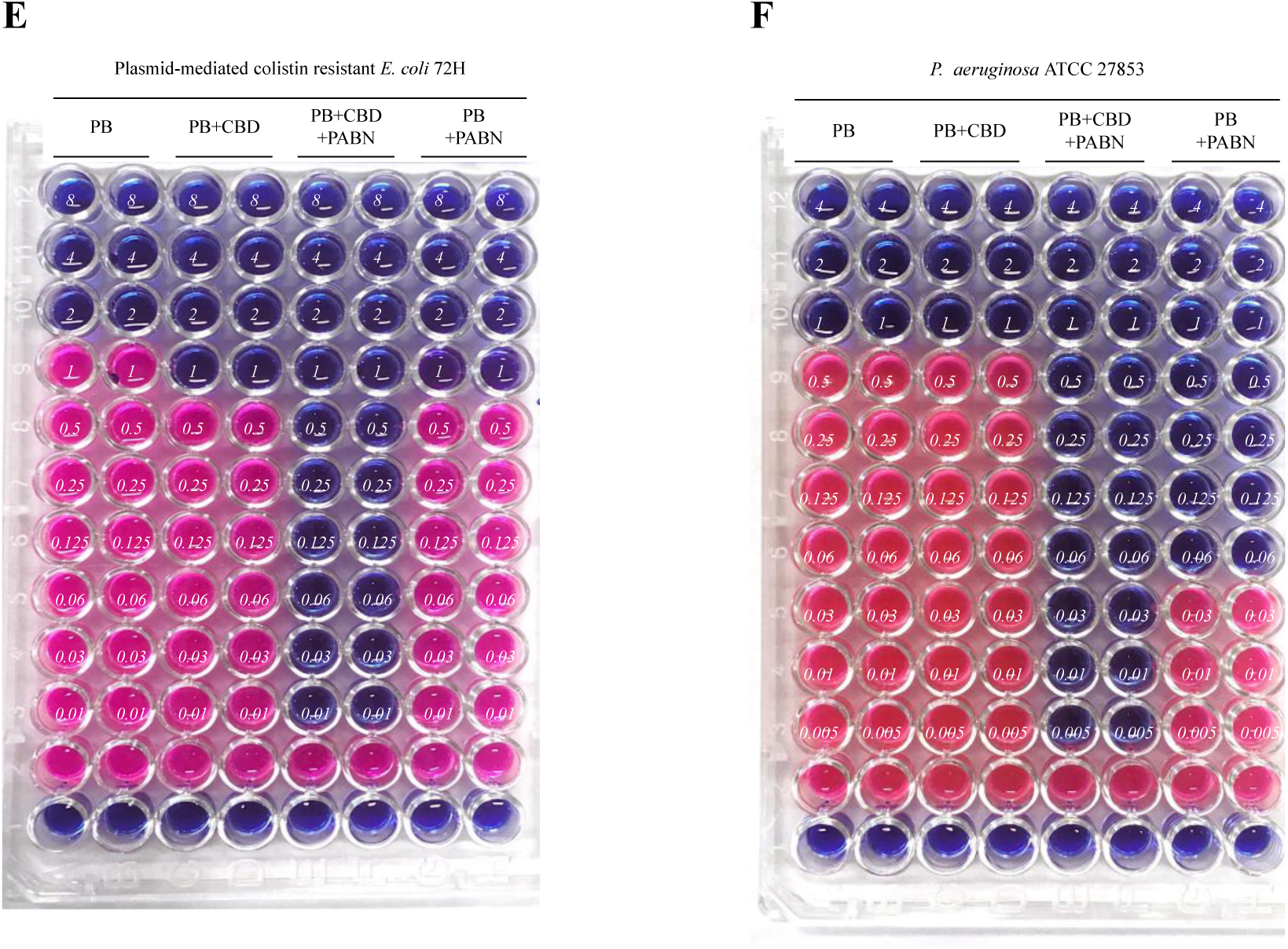
Representative screening assays results of antibacterial activity of the combination CBD + PB. The columns named “PB” are serial dilution of PB with no addition of CBD (PB MIC determination). The columns named “PB+CBD” are serial dilution of PB, plus a fixed concentration of CBD (256 µg/mL). The columns named “PB+CBD+PAβN” are serial dilution of PB, plus a fixed concentration of CBD (256 µg/mL) and a fixed concentration of PAβN (50 µg/mL). Finally, the columns named “PB+PAβN” are serial dilution of PB plus a fixed concentration of PAβN (50 µg/mL), with no addition of CBD. Blue wells show bacterial growth inhibition, while pink wells show bacterial growth. In each well, the numbers in italic refer to PB concentrations. Line “1 is MHB sterility control, while line “2” are bacterial growth control. Screening results of (**A**) PB-resistant *K. pneumoniae* C9, (**B**) PB-resistant *K. pneumoniae* L3 (left) and *K. pneumoniae* L8 (right), (**C**) *K. pneumoniae* L13 (left) and *K. pneumoniae* L14 (right), (**D**) *K. pneumoniae* L29 (left) and *K. pneumoniae* L18 (right), (**E**) Plasmid-mediated colistin resistant *E. coli* 72H (MCR-1), and (**F**) PB-susceptible *P. aeruginosa* ATCC 27853.

Regarding PB-resistant GNB (*Enterobacterales*), the combination of CBD plus low concentrations of PB (≤ 2 µg/mL) showed antibacterial activity against chromosomal PB- resistant GNB, including PB-resistant *K. pneumoniae* (Table 1 and Fig. 1A-D). However, for plasmid-mediated colistin-resistant (MCR-1) *E. coli* strains, we observed just a 1-fold reduction in the PB concentration compared to PB MIC (Table 1 and Fig. 1E).

The combination CBD + PB + PAβN showed antibacterial activity against GNB, so that lower PB concentrations were required when compared to the combination CBD + PB (Table 2 and Fig. 1). Interestingly, this combination (CBD + PB + PAβN) also showed antibacterial activity against *P. aeruginosa* and plasmid-mediated colistin-resistant (MCR-1) *E. coli* 72H strains (Table 2 and Fig. 1E-F).

**Table 2.**
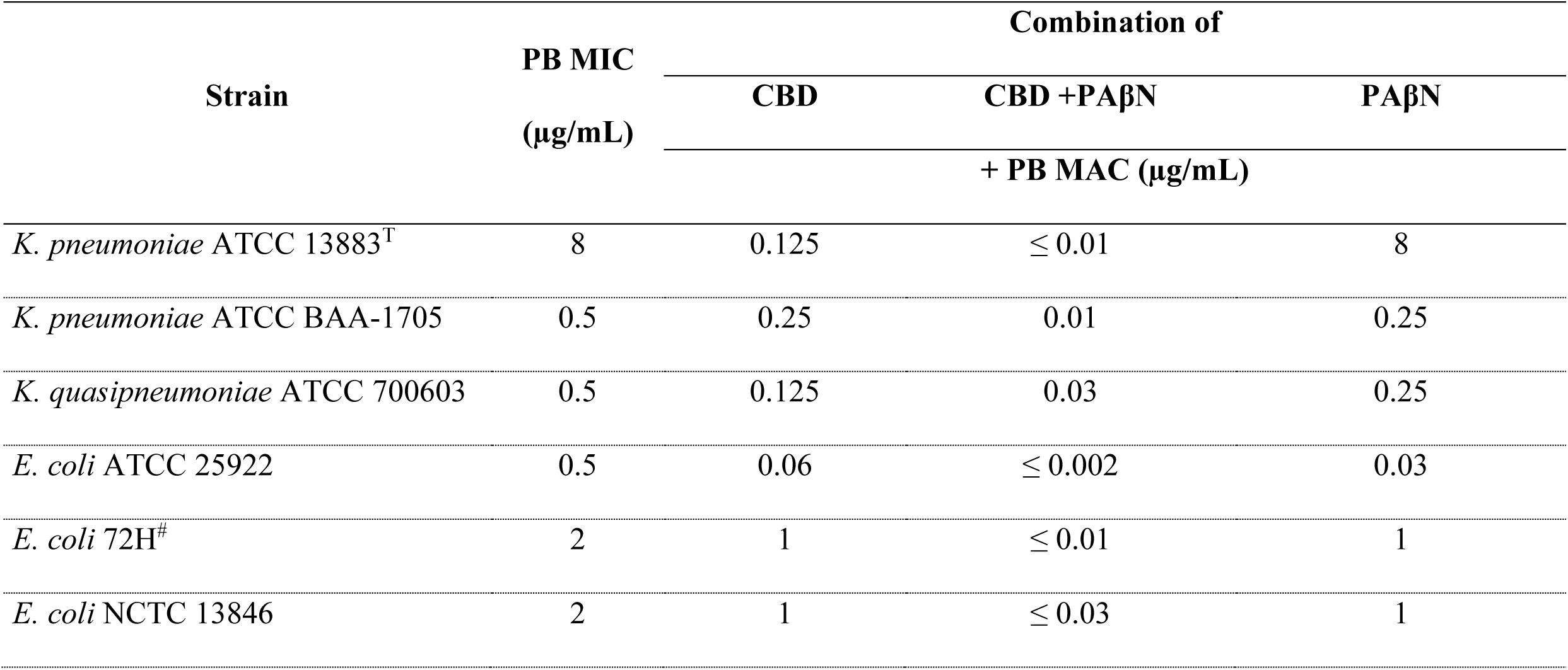
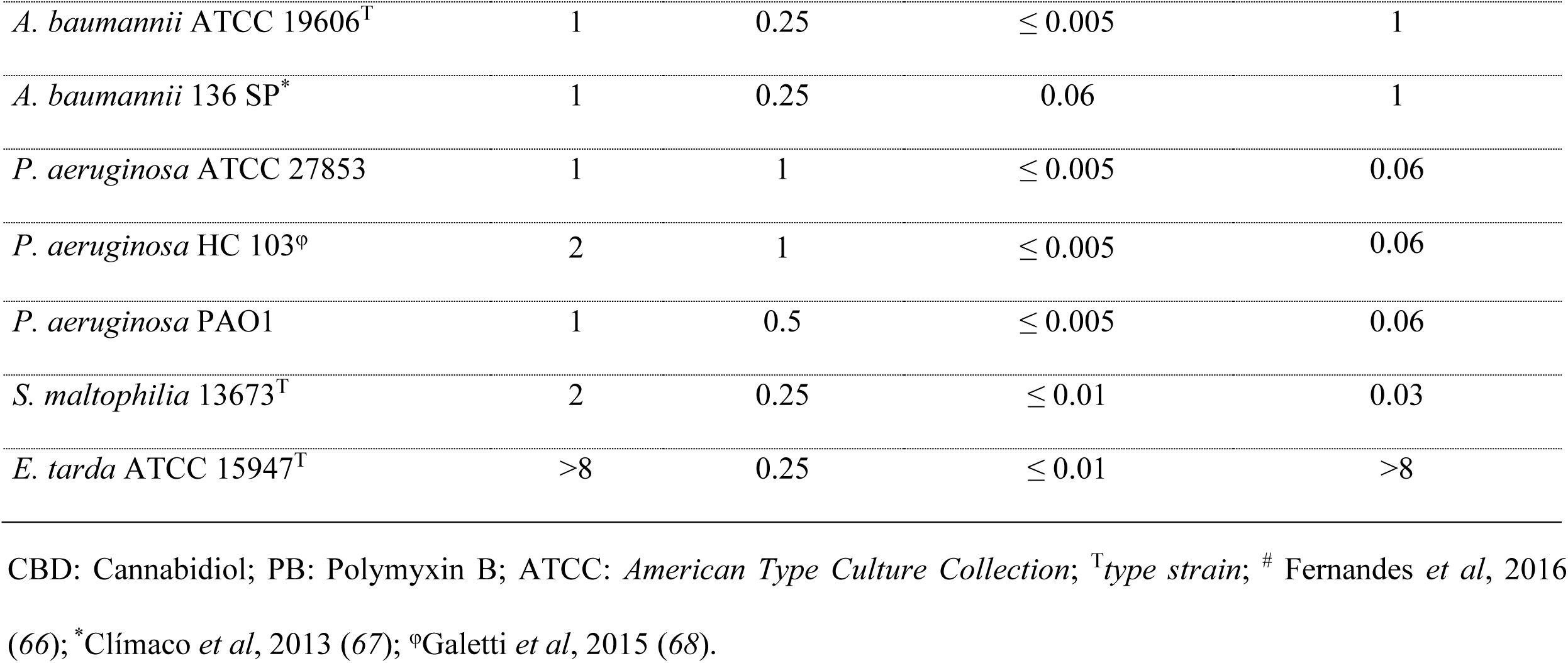
PB concentrations in the combination of CBD (256 µg/mL) + PB + PAβN (50 µg/mL) compared to PB concentrations in the combination of CBD (256 µg/mL) + PB. Lower PB concentrations were required in the presence of PAβN compared to the combination CBD + PB. This combination (CBD + PB + PAβN) also showed antibacterial activity against *P. aeruginosa* and plasmid-mediated colistin resistant *E. coli*.

Interestingly, CBD + PB also demonstrated antibacterial activity against intrinsically PB- resistant *Edwardsiella tarda* ATCC 15947 (Table 1). Nevertheless, for other intrinsically PB- resistant strains (*Burkholderia cepacia* ATCC 25416, *Morganella morganii* ATCC 8019, *Providencia rettgeri* ATCC 29944, *Proteus mirabilis* ATCC 29906, and *Serratia marcescens* subsp. *marcescens* ATCC 13880), no antibacterial activity of CBD + PB was observed.

Regarding intrinsically PB-resistant GNB, the combination of CBD + PB was not antibacterial even in the presence of PAβN. The exception was *E. tarda* ATCC 15947 for which the combination CBD + PB + PAβN was antibacterial, once again showing the antibacterial activity of CBD with lower PB concentrations (Table 2).

#### Checkerboard assay results

For BGN, the fractional inhibitory concentration index (FICI) of the combination CBD + PB was not calculated due to the absence of antibacterial activity (MIC) of CBD against GNB. Although, our results showed that for most of BGN (including PB-resistant *K. pneumoniae*) 4 µg/mL of CBD were enough to lead to bacterial growth inhibition when combined with low concentrations of PB (≤ 2 µg/mL) (Table 3 and Fig. 2A).

**Fig. 2.**
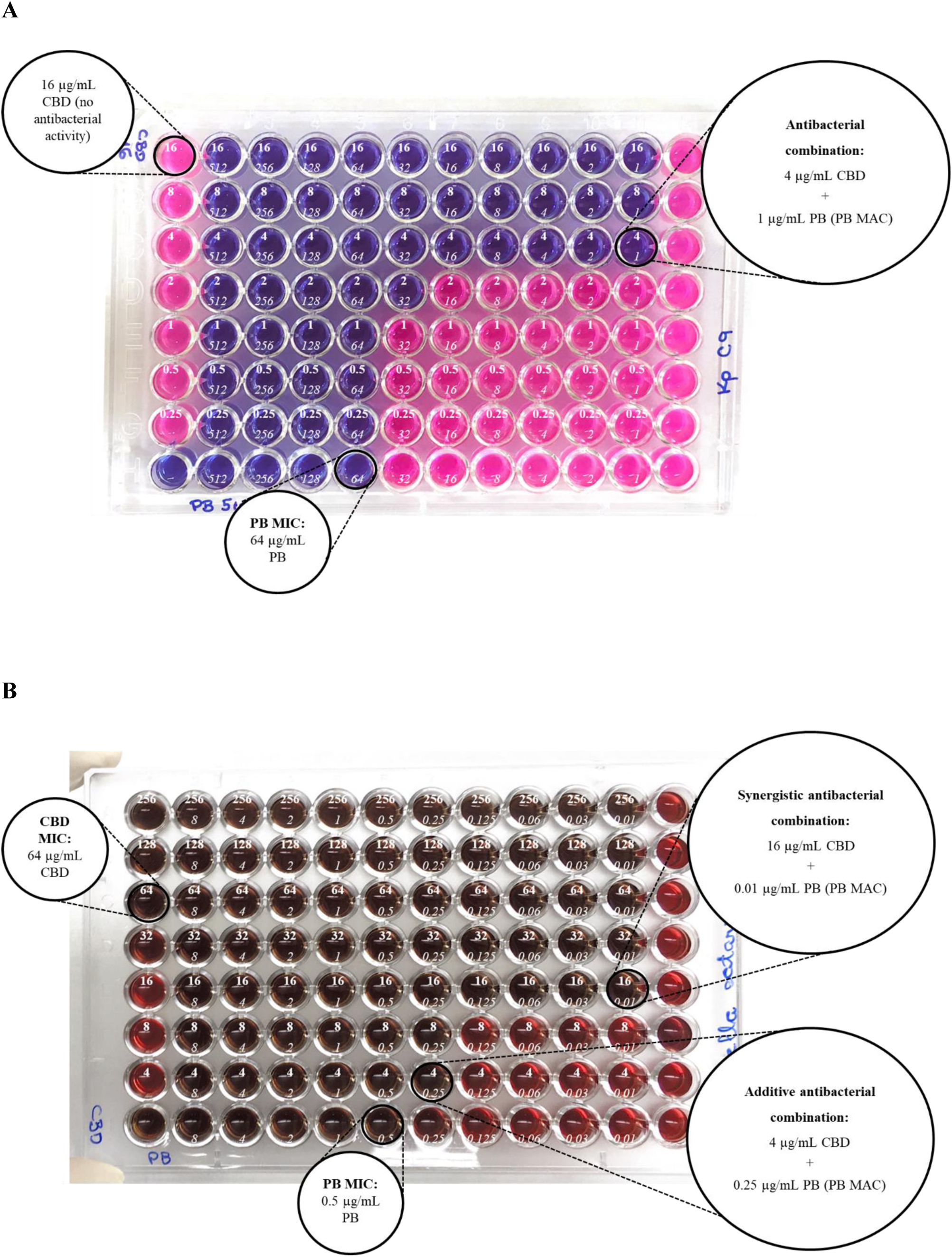
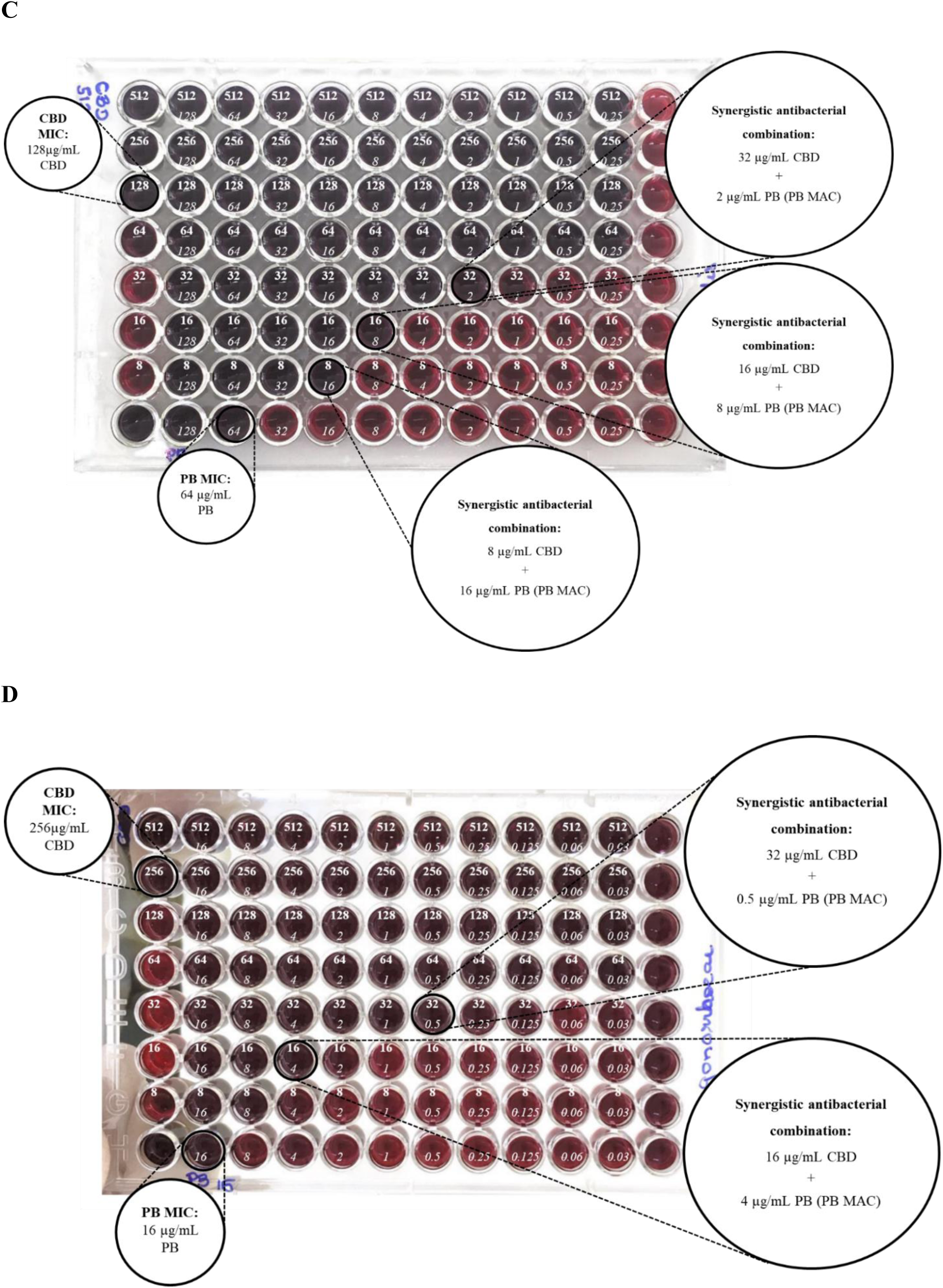
Representative checkerboard assays results showing the concentrations of CBD and PB (PB minimal antibiotic concentration [MAC]) required to the antibacterial activity of CBD in the combination CBD + PB. Blue wells are bacterial growth inhibition, while pink wells are bacterial growth. Serial dilution of CBD was performed horizontally (from the top to the bottom of the microplate), while serial dilution of PB was done vertically (from the left to the right of the microplate). In each well, the bold numbers of the top are referent to CBD concentrations, and the numbers in italic refer to PB concentration. Column “12” represents bacterial growth control, and the well “1H” is referent to the MHB sterility control. Checkerboard results for (**A**) PB-resistant *K. pneumoniae* C9, (**B**) *M. catarrhalis* ATCC 25238, (**C**) *N. meningitidis* ATCC 13077, and (**D**) *N. gonorrhoeae* ATCC 19424.

**Table 3.** Minimal concentrations of CBD and PB (PB minimal antibiotic concentration [MAC]) required to the antibacterial activity of the combination CBD + PB, according to checkerboard assay results. Plasmid-mediated colistin resistant *E. coli* 72H (MCR-1) and *P. aeruginosa* strains (ATCC 27853 and HC 103) were assessed in the presence of 50 µg/mL of PAβN.

| Strain | Antibacterial activity of the<br>combination CBD + PB (µg/mL) |  | PB MIC<br>(µg/mL) | PB MIC *versus*<br>PB MAC<br>(two-fold differences) |
| --- | --- | --- | --- | --- |
|  | CBD | PB MAC |  |  |
| *K. pneumoniae* ATCC 13883<sup>T</sup> | 4 | 0.03 | 8 | 8 |
| *K. pneumoniae* ATCC BAA-1705 | 4 | 0.25 | 0.5 | 1 |
| *K. pneumoniae* C9<sup>o</sup> | 4 | 2 | 64 | 5 |
| *K. pneumoniae* D1<sup>o</sup> | 4 | 0.25 | 1 | 2 |
| *K. pneumoniae* RP 62<sup>‡</sup> | 16 | 2 | 32 | 4 |
| *E. coli* ATCC 25922 | 4 | 0.06 | 0.5 | 3 |
| *E. coli* CTX-M-15 | 8 | 0.03 | 0.5 | 4 |
| *E. coli* 72H<sup>#</sup> | 4 + PAβN | 1 | 2 | 1 |
| *A. baumannii* ATCC 19606<sup>T</sup> | 4 | 0.125 | 1 | 3 |
| *A. baumannii* 136 SP<sup>*</sup> | 1 | 0.25 | 1 | 2 |
| *P. aeruginosa* ATCC 27853 | 4 + **PAβN** | 0.03 | 1 | 5 |
| *P. aeruginosa* HC 103<sup>¶</sup> | 4 + **PAβN** | 0.03 | 2 | 6 |

ATCC: American Type Culture Collection; <sup>T</sup>type strain; <sup>§</sup>Palmeiro *et al*, 2019 (64); <sup>‡</sup>Andrade *et al*, 2014 (65); <sup>#</sup>Fernandes *et al*, 2016 (66); <sup>\*</sup>Clímaco *et al*, 2013 (67); <sup>¶</sup>Galetti *et al*, 2015 (68).

For PB-susceptible *P. aeruginosa* ATCC 27853 and HC103 and plasmid-mediated colistin-resistant (MCR-1) *E. coli* 72H strains, the checkerboard assay was also performed in the presence of PAβN. The results showed that the combination CBD (4 µg/mL) + PB was antibacterial only in the presence of PAβN (Table 3).

For GND *M. catarrhalis* ATCC 25238, *N. meningitidis* ATCC 13077, and *N. gonorrhoeae* ATCC 19424, the FICI of the combination CBD + PB was calculated because both CBD and PB alone showed antibacterial activity (MIC). Thereby, CBD + PB showed additive and/or synergistic effect against these GND studied (Table 4 and Fig. 2B-D).

**Table 4.** Minimal concentrations of CBD and PB (PB minimal antibiotic concentration [MAC]) required to the antibacterial activity of the combination CBD + PB. Fractional Inhibitory Combination Index (FICI) values, according to checkerboard assay results. CBD showed antibacterial activity against GN diplococcus, so FICI was calculated and the effect of the combination was characterized.

| Strain | CBD MIC<br>(μg/mL) | Antibacterial activity of the combination |  | PB MIC<br>(μg/mL) | FICI | Effect |
| --- | --- | --- | --- | --- | --- | --- |
|  |  | CBD + PB (μg/mL) |  |  |  |  |
|  |  | CBD | PB MAC |  |  |  |
| *M. catarrhalis* ATCC 25238 | 64 | 4 | 0.25 | 0.5 | 0.56 | Additive |
|  |  | 16 | 0.01 |  | 0.27 | Synergic |
| *N. meningitidis* ATCC 13077 | 128 | 32 | 2 | 64 | 0.28 | Synergic |
|  |  | 16 | 8 |  | 0.25 |  |
|  |  | 8 | 16 |  | 0.31 |  |
| *N. gonorrhoeae* ATCC 19424 | 256 | 32 | 0.5 | 16 | 0.155 | Synergic |
|  |  | 16 | 4 |  | 0.31 |  |

## DISCUSSION

Our main contributions were:

(i) CBD antibacterial activity data against a broad panel of GN bacteria (30 different species; 73 strains), comprehending type-strains, standard strains, and clinical isolates (including international high-risk clones, MDR and XDR strains, and also susceptible strains);
(ii) Comprehensive interpretation of CBD antibacterial activity against LOS-expressing GND (*N. meningitidis*, *N. gonorrhoeae*, and *M. catarrhalis*) compared to LPS-expressing GNB;
(iii) Antibacterial activity data of the combination CBD + PB against intrinsic PB-resistant, and chromosomal- and plasmid-acquired PB-resistant GNB, highlighting PB-resistant *K. pneumoniae*;
(iv) Comprehensive interpretation of the biological effect of the combination CBD + PB (+ PAβN);
(v) Discussion about pharmacokinetic perspectives of CBD repurposing as antibacterial agent: CBD alone against GP bacteria or of the combination CBD + PB against GN bacteria, highlighting PB-resistant GNB.

### CBD antibacterial activity against GP bacteria, *M. tuberculosis*, and LOS-expressing GND

We observed antibacterial activity of ultrapure CBD against GP bacteria, *M. tuberculosis*, and LOS-expressing GND; but not against GNB, as recently described (*17–19*). Preliminary results regarding CBD activity in GP bacteria and its absence in GN bacteria (including MDR and XDR strains) were partially presented by us at ASM Microbe 2018 and published in the abstract book (*15*).

Our results additionally provide CBD antibacterial activity data against other species: *Enterococcus casseliflavus, Staphylococcus lugdunensis, Micrococcus luteus,* and *Rhodococcus equi*, and also against different phenotype/genotype of GP bacteria (Table S1). In general, our data and previous reports showed CBD MICs ranging from 2 to 4 µg/mL against GP bacteria, including vancomycin-resistant *E. faecium* (VRE), and methicillin-resistant (MRSA) and vancomycin-intermediate resistant (VISA) *S. aureus*, all of them listed in the WHO priority pathogens list for R&D of new antibiotics.

Our data differ from data by Blaskovich *et al*. (2021) about CBD MICs against *S. pneumoniae, S. pyogenes, N. meningitidis, N. gonorrhoeae, M. catarrhalis,* and *M. tuberculosis* (*19*). Differences in the MIC values observed may be related to the different material and methods used in each one of the studies. For fastidious bacteria, such as *S. pneumoniae, S. pyogenes, M. catarrhalis,* and *N. meningitidis*, we used MH-F broth (CAMHB supplemented with 5% lysed horse blood + 0.1% β-NAD 20 mg/mL), according to standard protocol by EUCAST (*24*). For *N. gonorrhoeae*, we performed the broth microdilution method instead of agar dilution. Blaskovich et al. (2021) used a broth culture medium composed by a lower percentage of lysed horse blood (3%) for *S. pneumoniae* and *S. pyogenes*, and a modified broth according to “American Type Culture Collection (ATCC)” for *Neisseria* spp., which does not contain blood (*19*). The presence of blood on culture medium composition (e.g., MH-F broth) increases CBD MIC, as observed in our results for *S. aureus* (4-fold dilution increased CBD MIC) when compared to only CAMHB as well as previously described (*14*).

Besides, we used only non-treated plates (non-binding surface), as recommended for the broth microdilution method, according to standard protocols from EUCAST / Clinical & Laboratory Standards Institute (CLSI) (*22, 23*). These differences may have contributed to our CBD MIC values against fastidious bacteria be higher than the values presented by Blaskovich et al. (2021) (21).

Regarding to *M. tuberculosis*, our results showed lower CBD MIC values than Blaskovich et al. (2021), but some experimental variations may also explain the differences. Blaskovich et al. (2021) used a period of incubation of 5 days, the addition of 12.5uL of 20% tween 80 into resazurin, and culture medium supplemented with ADC (albumin, dextrose, catalase) (Difco Laboratories), 0.5% glycerol, and 0.02% tyloxapol. We used a period of incubation of 7 days, no addition of tween 80 into resazurin, and culture medium supplemented with OADC (oleic acid, albumin, dextrose, catalase) and glycerol 0,4%. The difference between OADC and ADC is the presence of oleic acid at 0.5g/L.

### CBD antibacterial activity against GN bacteria (GNB and GND)

Our results reinforce that CBD alone is not antibacterial against GNB (MDR or susceptible to antibiotics), since we evaluated 27 species (70 strains) of the most GNB species involved in healthcare-associated infections (HAI) as well as in community infections, expanding the panel of GNB and human pathogens investigated: *Klebsiella oxytoca, Klebsiella aerogenes, Pantoea agglomerans, Cronobacter sakazakii, Providencia rettgeri, Salmonella enterica* subsp*. enteric* (serovar Enteritidis)*, Shigella flexneri, Plesiomonas shigelloides, Hafnia alvei, Edwardsiella tarda, Yersinia enterocolitica* subsp*. Enterocolitica, Pseudomonas putida, Aeromonas hydrophila,* and *Alcaligenes faecalis* subsp*. faecalis* (Table S1).

Our data also showed no role of efflux pumps commonly involved in antibiotic extrusion from GN cell. Thereby, we hypothesize that CBD could not be active due to low permeability through cell envelope (outer membrane) of GNB.

The absence of antibacterial activity of CBD against GNB may be related to LPS molecules and outer membrane proteins, from outer membrane, which would lead to the impermeability of macromolecules and limited diffusion of hydrophobic molecules, such as CBD (*17–19*). Our results of *in vitro* CBD antibacterial activity against GP bacteria and *M. tuberculosis*, and the absence of CBD antibacterial activity against GNB, enhance LPS role in hinder CBD activity.

The CBD hydrophobic chemical structure may suggest the interaction of the molecule with lipid membranes, as described by Guard *et al* (2020) for eukaryotic cells, altering the biophysical properties of the membrane and affect the metabolism of lipids and cholesterol (*11, 25*). The bacterial membrane was also suggested as a possible bacterial target for cannabinoids (cannabigerol [CBG] and CBD) (*13, 17, 19*). Also, Blaskovich *et al*. (2021) also showed that CBD bactericidal concentrations against *S. aureus* lead to inhibition of protein synthesis, DNA, RNA, and peptidoglycan (*19*). Nevertheless, CBD specific antibacterial mechanism(s) of action was not yet fully elucidated.

Furthermore, we have evaluated bacteria that express LOS, which lack the O-antigen (*N. meningitidis*, *N. gonorrhoeae,* and *M. catarrhalis*), on their outer membrane, instead of LPS (*4, 26*). Thereby, the antibacterial activity of CBD against these bacteria could suggest O-antigen specific role in preventing CBD antibacterial activity (possibly due to the steric effect, hindering CBD to reach its molecular target).

Even considering that *A. baumannii* and *H. influenzae* also have LOS molecules on their external membrane, the core polysaccharide of these bacteria presents a different sugar composition (*4, 27*). This fact could explain the absence of CBD antibacterial activity against these GN bacteria. CBD and CBG have antibacterial activity against *A. baumannii* only in the absence of whole LOS, according to previous studies (*17, 19*).

Therefore, our results contribute to a better understanding about CBD antibacterial mechanism(s) of action, which could guide future studies.

### Antibacterial activity of the combination CBD + PB

Previous studies reported the antibacterial activity of CBD in combination with PB, although only few bacterial species and strains were evaluated (*E. coli*, *P. aeruginosa*, *K. pneumoniae* and *A. baumannii*) (*17, 19*). We partially published preliminary results regarding the antibacterial activity of the combination CBD + PB (including carbapenem-resistant, chromosomal- and plasmid-acquired PB-resistant, and intrinsic PB-resistant GNB) in the abstract book of the 30th European Congress of Clinical Microbiology & Infectious Diseases (ECCMID 2020) (*28*).

Our data provide additional results of the effect of the combination CBD + PB against several other GN bacteria, including GNB and GND (11 species, 51 strains), highlighting the critical priority pathogens: carbapenem-resistant (CRE; e.g., carbapenemase producers) and 3^rd^- 4^th^ generation cephalosporin-resistant (e.g., extended-spectrum beta-lactamase [ESBL] producers) *Enterobacterales* (e.g., *K. pneumoniae* and *E. coli*), carbapenem-resistant *A. baumannii* (CRAB) and carbapenem-resistant *P. aeruginosa*, from WHO priority pathogens list for R&D of new antibiotics. These strains include MDR and XDR international high-risk clones (e.g., *K. pneumoniae* ST 258 and ST 11, *E. coli* ST 131, *P. aeruginosa* ST 277, and *A. baumannii* ST 109) (Table 1).

Since the antibacterial activity of the combination CBD + PB was observed against GNB, these results point to the existence of a CBD molecular target also in GN bacteria and indicate that its exposure is dependent on outer membrane disruption or destabilization for antibacterial activity.

For the most GNB, our checkerboard results showed that 4 µg/mL of CBD was the minimal concentration needed to the antibacterial activity into combination CBD + PB. Indeed, PB concentrations ranging of one up to eight two-fold lower than PB MIC were needed to the antibacterial activity of the combination CBD + PB. For PB-resistant GNB (including *K. pneumoniae* and plasmid-mediated colistin resistant [MCR-1] *E. coli* strains), 4 µg/mL of CBD were enough to lead to bacterial growth inhibition when combined with clinically relevant PB concentrations (PB MIC ≤ 2 µg/mL) (Table 3 and Fig. 2A).

Among intrinsic PB-resistant GNB, the combination of CBD + PB showed antibacterial activity only against *E. tarda*. These results may be related to the these bacteria’s different intrinsic resistance mechanisms, involving different molecular pathways from two-component systems (*2, 29*).

For GND *N. meningitidis*, *N. gonorrhoeae,* and *M. catarrhalis*, FICI calculation demonstrated additive or synergistic effects for the combination CBD + PB (Table 4 and Fig. 2B-D). Nevertheless, PB is not used for the treatment of infections caused by these GND; however, the synergistic effect may suggest a new insight for this bactericidal activity of the combination CBD + PB, highlighting that PB also neutralizes the endotoxin Lipid A of the LPS/LOS of GN bacteria (*30*).

### Biological effect of the combination CBD + PB (+ PAβN)

Our results showed that low PB concentrations (lower than PB MIC against each GNB evaluated) are enough to lead to the minimal outer membrane disruption (or destabilization) required to allow antibacterial activity of CBD in GNB (except *P. aeruginosa* and plasmid-mediated colistin-resistant [MCR-1] *E. coli* strains). Thus, this low PB concentration could be considered the minimal antibiotic concentration (MAC), nominated as the minimal concentration that produces any biological effect on bacterial cells (e.g., outer membrane destabilization) (*31*).

In this context, the lower fold reduction among PB-susceptible *K. pneumoniae* strains compared to PB-resistant *K. pneumoniae* strains could be explained by a short-range between PB MAC and PB MIC for these bacteria, because the concentration required to the outer membrane destabilization is similar among PB-susceptible and PB-resistant strains. Thus, as PB MIC is higher for PB-resistant strains, the range between PB MAC and PB MIC is more extensive among these strains, so it is possible to observe a higher number of fold reduction in the PB concentration (into combination CBD + PB) compared to PB MIC.

Based on our results as well as explanation, we believe that: CBD does not decrease the PB MIC against bacteria, since “MIC” is the “minimum inhibitory concentration” of only one antibacterial agent, so, in combination with another substance, the concept of MIC reduction is mistaken and should not be used. The lower PB concentrations in the combination CBD + PB are the PB MAC.

Similarly, CBD does not restore PB susceptibility in PB-resistant strains, because CBD is the antibacterial agent in the combination CBD + PB, considering that PB concentrations are lower than PB MIC (so, it is subinhibitory) against these PB-resistant GN bacteria.

Furthermore, once CBD MIC could not be determined for GNB, the antibacterial activity of the combination CBD + PB could not be categorized as synergistic or additive, based on checkerboard assay, due to lack of mathematical factor (MIC) to calculate the FICI. Thereby, to attribute the maximum concentration evaluated in the experiments like a “MIC” is also a mistake.

The combination CBD + PB + PAβN was effective also against *P. aeruginosa* and plasmid-mediated colistin resistant (MCR-1) *E. coli* strains, for which CBD + PB does not have activity. Although, these results could not be related to efflux inhibition by PAβN *per se*, because CBD antibacterial activity alone was not detected in the presence of PAβN. Thereby, our results suggest PAβN permeabilization of the outer membrane contributing to CBD activity, as similarly described for β-lactams in the presence of PAβN against *P. aeruginosa*, or sensitization of *P. aeruginosa* to antibiotics (e.g., vancomycin) that are typically incapable of crossing the outer membrane (*32*). Indeed, combination CBD + PB in the presence of PAβN decreases considerably the PB concentrations necessary to allow CBD activity.

PAβN is a substance commonly used for bacterial efflux pump inhibition in *in vitro* assays. Although, it is not currently used as a drug in clinical practice, there is a suggestion of potential use as an antibiotic adjuvant that might reduce the effective doses of current drugs which need increased outer membrane permeability (*32*).

### Pharmacokinetic perspectives of CBD repurposing as antibacterial

In 2018, the Food and Drug Administration (FDA, United States) approved CBD (Epidiolex^®^, Greenwich Biosciences, Inc.) for the treatment of patients who have Lennox-Gastaut syndrome or Dravet syndrome (*33*). Likewise, according to European Medicines Agency, Epidyolex^®^ (GW Pharma [International] B.V) was authorized in the European Union in 2019, for the same therapeutic indication when in addition to clobazam, another anti-epileptic drug (*34, 35*). In 2020, the Brazilian Health Regulatory Agency (ANVISA) approved CBD (Canabidiol Prati-Donaduzzi^®^, Toledo, PR, Brazil) for the treatment of pharmacoresistant epilepsy or refractory epilepsy (*36*).

According to the Biopharmaceutics Drug Disposition Classification System, CBD is a class 2 drug, showing low water solubility and high permeability/metabolism depending on CYP3A4, CYP2C19, UGT1A9 and UGT2B7 enzymes (*37–41*).

The clinical CBD pharmacokinetics studies following oral administration were evaluated in different formulations such as capsules, solutions and oromucosal preparations (*42, 43*). However, the oral bioavailability of CBD is low (6-25%), highly variable and increased with coadministration of a high-fat meal (*42, 44, 45*). Additionally, CBD bioavailability (59%) could be higher through pulmonary administration (*46*).

The antibacterial efficacy of CBD would be dependent on CBD unbound fraction be able to achieve the infection target site. CBD unbound fraction in healthy volunteers is 6.98% and increases to 11.69% in severe hepatic impairment (*45*).

Intravenous (IV) administration is the most usual administration route of antimicrobial therapy in critically ill patients, and CBD pharmacokinetics studies following IV administration have been studied in some clinical trials (*43, 47, 48*).

CBD pharmacokinetics after a 20 mg IV dose was evaluated by a previous study, which showed clearance values of approximately 80 L/h and volume of distribution of 52 L in 70 kg individuals (45). Thus, as an exercise of translational pharmacokinetics could be done using a classic equation and considering linear pharmacokinetics:

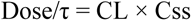

Where τ represents the dosing interval, CL is the total clearance, and Css represents the steady-state mean plasma concentration.

For the antibacterial activity of CBD, an IV administration of CBD doses of approximately 2 g/12 h or 4 g/12 h would result in plasma exposure higher than, respectively, 2 µg/mL or 4 µg/mL, that are CBD MIC against several GPC (e.g., *Enterococcus* spp. and *Staphylococcus* spp.) (Table S1).

For the antibacterial activity of the combination CBD + PB against the most PB-resistant GNB, especially *K. pneumoniae*, 4 µg/mL of CBD seems to be the better concentration of CBD for antibacterial activity in the combination with ≤2 µg/mL of PB (PB MAC) (Table 3 and Fig. 2A). PB MIC ≤2 µg/mL against the GNB *Enterobacterales* (e.g., *K. pneumoniae* and *E. coli*)*, P. aeruginosa* and *A. baumannii* categorizes theses bacteria as susceptible strain; and there is a high likelihood of therapeutic success using a standard dosing regimen of PB; safe and effective as ‘last-resort’ antibiotic to treat severe infections due to MDR GNB (*22, 49*).

To date, there are no published studies evaluating the administration of doses as high as those discussed above. However, considering that CBD shows low water solubility (12.6 mg/L) and high lipophilicity (logP of 6.3), this present a limitation for the development of an IV pharmaceutical formulation compatible with the doses proposed here (*50*). Nevertheless, a study presented a simple formulation of both CBD and delta-9-tetrahydrocannabinol (THC) 1.6 mg IV, which was administered in eight healthy individuals (*46*).

The optimization of the CBD IV administration and pharmacokinetic parameters could be achieved by nanomaterial-based strategies, using nanocarriers to increase the solubility, stability and efficacy of IV CBD for antibacterial therapy (*51*).

Regarding CBD safety and tolerability, one study has evaluated the adverse effects following IV administration, while other clinical trials have evaluated its use in single or multiple oral doses (*52*). Bhattacharyya *et al*. (2010) performed several behavioral tests in healthy volunteers after the IV administration of CBD (5 mg) concomitantly or not with THC. Besides the block of psychotic symptoms from THC by CBD, the authors have not discussed the safety concerns of the IV administration of CBD (*52*). Meyer *et al*. (2018) evaluated a CBD- THC dose of 1.6 mg administered IV and pulmonary and described that adverse reactions presented by the administration of both drugs were mainly psychological (such as the feelings of relaxation, concentration, and somnolence), while psychotic symptoms presented such as confusions and changes in perception were mainly associated as typically THC related. (*46*).

The therapeutic oral dose of 1.5 g CBD per day was reported to be well tolerated and safe in humans (*53–55*). A recent randomized, double-blind, and placebo-controlled clinical study evaluated adverse events in two different arms: a single oral dose of CBD up to 6 g and multiple doses of 0,75 g to 1.5 g twice daily for 7 days. Both arms presented only mild or moderate adverse events, being the most frequent diarrhea, nausea, headache, and somnolence. Additionally, the safety of a single 1.5 g CBD dose was investigated in healthy volunteers fasted or following a high-fat meal. More subjects experienced adverse events following a high-fat meal, which was explained by the enhanced bioavailability. Nevertheless, the authors concluded no safety concerns (*45*).

PB standard dosage is administered IV and is predominantly cleared by unknown nonrenal pathways (*56*). So, it is not yet possible to predict CBD + PB interactions considering that there is no data regarding the involvement of metabolism enzymes and/or drug transporters in PB disposition.

Our *in vitro* results demonstrated that the combination CBD + PB is a potential alternative for clinical use as an antimicrobial treatment in MDR bacterial infection, including PB-resistant GNB. Future studies should evaluate the PK/PD relationship of the combination CBD + PB, highlighting the potential PB dose reduction when combined, which would allow to use PB even in therapeutical low concentrations, decreasing dosage of this antibiotic and mitigating its neurotoxicity and nephrotoxicity.

Several approaches have been used to better understand and more efficiently combat bacterial resistance to antibiotics, mainly against superbugs (e.g., PB-resistant Gram-negative bacilli) (*57, 58*). Drug repurposing may represent a faster approach to identify new antimicrobials, since preclinical and clinical parameters of these drugs are already established (*59–61*). Thereby, CBD antibacterial activity might be considered for drug repurposing and evaluated in clinical studies (e.g., expanded access) initially against immediately life-threatening condition or serious infections (*62, 63*).

CBD antibacterial activity alone could be evaluated against:

(i) XDR VISA (also optimizing pharmacokinetics/pharmacodynamics [PK/PD] parameters of clinical use of vancomycin);
(ii) XDR *M. tuberculosis* (also using in combination with alternative tuberculostatic).

Moreover, the antibacterial activity of the combination CBD + PB might also be investigated against infection due to PB-resistant GNB, which other options of antibiotics are ineffective:

(i) CRE (e.g., *Klebsiella pneumoniae carbapenemase* [KPC] producers), ceftazidime-avibactam (CAZ-AVI)-resistant, and XDR strains;
(ii) CRAB (for which CAZ-AVI is ineffective), and XDR strains;
(iii) Pandrug-resistant (PDR) strains.

### Final considerations

In conclusion, our *in vitro* assays showed CBD antibacterial activity against different GP bacterial species, LOS-expressing GND, and *M. tuberculosis*, contributing to new insights regarding CBD antibacterial activity in future studies. We demonstrated the antibacterial activity of the combination CBD + PB against PB-resistant GNB, as well as additive/synergistic effect against LOS-expressing GND. Our results show translational potential and CBD should be further explored as a potential antibacterial agent by clinical trials. The antibacterial efficacy of the combination CBD + PB against MDR and XDR GNB, highlighting PB-resistant *K. pneumoniae*, is particularly promising.

## MATERIALS AND METHODS

### Study design

CBD antibacterial activity was evaluated against a broad diversity of bacteria (43 different species, 94 strains), including GN bacteria (comprehending different cell wall composition and structure), GP bacteria, and *M. tuberculosis*, using the broth microdilution method. The antibacterial activity of the combination CBD + PB was also investigated for GN bacteria (11 species, 51 strains) using the broth microdilution method, as well as *checkerboard* assay (7 species, 15 strains), mainly against intrinsic PB-resistant, and chromosomal- and plasmid-acquired PB-resistant GNB. Furthermore, we present a discussion about pharmacokinetic perspectives of the combination of CBD + PB against PB-resistant GNB.

### Investigation of CBD antibacterial activity against GP and GN bacteria

Antibacterial activity of CBD was investigated against a broad panel of different bacterial species, comprehending GP (13 different species; 21 strains), GN (30 different species; 73 strains) bacteria, and *M. tuberculosis* (2 strains), including type-strains, quality control strains, and clinical isolates (MDR and XDR strains, international high-risk clones, and also susceptible strains) (Table S1) (*64–68*).

The reference broth microdilution method was used to determine CBD MIC, according to standard procedures (EUCAST/CLSI) (*23, 24*). Cation-Adjusted Mueller Hinton II Broth (CAMHB) (BBL™, Becton Dickinson), untreated polystyrene flat-bottom 96-wells microplates, and ultrapure CBD (99.6%; BSPG-Pharm, Sandwich, UK) were used. Methanol (Sigma-Aldrich) was used as CBD solvent, ranging from 0.006 to 327.68 µL/mL on CAMHB. Our previous standardization showed that methanol does not antibacterial in these concentrations.

For fastidious bacteria, such as *Streptococcus* spp*., Neisseria* spp*., Moraxella catarrhalis,* and *Haemophilus influenzae*, Cation-Adjusted Mueller Hinton II Broth (CAMHB) (BBL™, Becton Dickinson) supplemented with defibrinated horse blood and β-NAD, named MH-F, was used as recommended by EUCAST (*24*).

Two-fold serial dilution (256 - 0.5 µg/mL) of CBD were initially evaluated and MIC values were determined as the lowest concentrations of CBD that inhibit visible bacterial growth in broth culture medium. Polymyxin B and vancomycin were used as controls for GN and GP bacteria, respectively. In addition, ciprofloxacin was used as control for GND (*Neisseria* spp*., M. catarrhalis)*, and ampicillin for *S. pneumoniae* and *H. influenzae* (*22*). The assays were done in technical and experimental replicates.

Beyond visual evaluation of growth inhibition, 30 µL of 0.01 to 0.02% aqueous solution of resazurin sodium salt (Sigma-Aldrich) were added in each well of the microplate. After 30-60 minutes for GNB and GPC, and 60-120 minutes for GND and *Enterococcus* species., the metabolic activity and proliferation of bacterial cells was visually determined by bioreduction of the dye (blue), which is converted into resorufin (pink) in the presence of viable bacteria (*69*). This colorimetric step was additionally performed with the objective of better visualization of MIC.

CBD MBC was also verified for GPC: *Enterococcus faecalis* (ATCC 29212 and ATCC 51299), *E. faecium* (NCTC 7171 and ATCC 51559), and *S. aureus* (ATCC 29213 and ATCC 700699); as well as to GND: *N. meningitidis* ATCC 13077, *N. gonorrhoeae* ATCC 19424, and *M. catarrhalis* ATCC 25238. MBC method was performed after visual evaluation of growth inhibition by subculturing onto Mueller Hinton Agar or Mueller Hinton Agar with Blood (Difco™, Becton Dickinson) plates in the absence of CBD. MBC values were determined as lowest concentration of CBD that prevents bacterial colony-forming unit’s growth in solid culture medium.

### Investigation of CBD antibacterial activity against *M. tuberculosis*

We used the reference broth microdilution method to determine CBD MIC against *M. tuberculosis* H37Rv (ATCC 27294), and also against rifampicin and isoniazid-resistant *M. tuberculosis* CF86 (MDR clinical isolate), according to the standard procedures (*70*). Middlebrook 7H9 broth (Sigma-Aldrich) supplemented with 10% OADC (oleic acid, albumin, dextrose, catalase) and glycerol 0,4% was used and two-fold serial dilution (256 - 1 µg/mL) of CBD were evaluated. Rifampicin and isoniazid were used as control (1 – 0.004 µg/mL). Each plate was incubated for seven days, under 37°C and 5% CO_2_. After incubation, 30 µL of 0.01% aqueous solution of resazurin sodium salt (Sigma-Aldrich) were added in each well of the microplate, and 24 hours later MIC was determined through fluorescence reading (530/590 nm). The assay was performed in two independent experiments (*71*).

### Investigation of CBD antibacterial activity against GNB in the presence of efflux pump inhibitors

Efflux inhibition assay was performed to investigate CBD extrusion through efflux pumps from susceptible and resistant Gram-negative ESKAPE pathogens (*K. pneumoniae, A. baumannii, P. aeruginosa, Enterobacter* species) as well as *E. coli* and *S. maltophilia* strains.

Broth microdilution method to determine CBD MIC was performed in the presence and in the absence of PAβN (Sigma-Aldrich) (50 µg/mL), reserpine (Sigma-Aldrich) (50 µg/mL), and curcumin (256 µg/mL), in different assays (*72, 73*). We considered that a minimal 3-fold reduction in the MIC values in the presence of efflux pump inhibitors would be suggestive of efflux-mediated resistance.

### Investigation of the antibacterial activity of CBD in combination with PB against GN bacteria

#### Broth Microdilution method with fixed concentration of CBD (256 µg/mL)

For GN bacteria (11 species, 51 strains), antibacterial activity of the combination CBD + PB was evaluated against PB-susceptible and PB-resistant bacteria, including acquired (chromosomal and plasmid-mediated colistin-resistant [MCR-1] *E. coli*) (Table 1), and intrinsically-resistant GNB (*B. cepacia* ATCC 25416, *M. morganii* ATCC 8019, *P. rettgeri* ATCC 29944, *P. mirabilis* ATCC 29906, and *S. marcescens* subsp. *marcescens* ATCC 13880).

To investigate the antibacterial activity of the combination CBD + PB, initially a screening was performed using broth microdilution method, with adaptations: Two-fold serial dilution (512 - 0.02 µg/mL) of PB was evaluated in the presence of 256 µg/mL of CBD (fixed concentration) in each well, including the bacterial growth control wells.

Furthermore, the antibacterial activity of the combination CBD (256 µg/mL of fixed concentration) + PB (two-fold dilution, 256 – 0.005 µg/mL) was also evaluated in the presence of PAβN (50 µg/mL), also including the bacterial growth control wells.

#### Checkerboard assay

For *K. pneumoniae* (n=5), *E. coli* (n=3), *A. baumannii* (n=2), *P. aeruginosa* (n=2), *N. meningitidis* (n=1), *N. gonorrhoeae* (n=1), and *M. catarrhalis* (n=1), the checkerboard assay was performed to assess the *in vitro* antibacterial activity of the combination CBD + PB, in untreated polystyrene 96-wells microplates, as previously described (*74*). Final PB concentrations ranged from 0.01 to 512 µg/mL and CBD concentrations ranged from 4 to 256 µg/mL. For *P. aeruginosa* and plasmid-mediated colistin resistant (MCR-1) *E. coli* strains, the assay was performed also in the presence of 50 µg/mL of PAβN.

The checkerboard assay results were analyzed considering the fractional inhibitory concentration index (FICI) that categorize the combination (of two different substances, e.g., CBD + PB) as synergistic, additive, indifferent, or antagonist. FICI calculation of the antibacterial activity of the combination CBD + PB considered the best well(s) where the concentrations of PB (closest to 2 µg/mL or lower) combined to CBD (lowest concentrations) inhibited bacterial growth into the combination CBD + PB. Thereby, FICI was calculated as: FICI = (CBD concentration in combination / MIC_CBD_) + (PB concentration in combination / MIC_PB_). FICI values were interpreted as: ≤0.5 (synergy), >0.5-1 (additive), 1-4 (indifference) and >4 (antagonism) (*74*).

For “broth Microdilution method with fixed concentration of CBD (256 µg/mL)” as well as “checkerboard assay”, beyond visual evaluation of growth inhibition, 30 µL of 0.01 to 0.02% aqueous solution of resazurin sodium salt (Sigma-Aldrich) were added in each well of the microplate. After 30-60 minutes for GNB, and 60-120 minutes for GND, the metabolic activity and proliferation of bacterial cells was visually determined by bioreduction of the dye (blue), which is converted into resorufin (pink) in the presence of viable bacteria (*69*). This colorimetric step was additionally performed with the objective of better visualization of MIC.

## Supplementary Materials

Fig. S1. CBD MIC results against *S. aureus* ATCC 29213 observed from broth microdilution assay using MH-F broth and using CAMHB.

Fig. S2. Representative screening assays results of antibacterial activity of the combination CBD + PB against PB-susceptible *K. pneumoniae* BAA-1705. Table S1. CBD MIC results against bacterial strains studied.

## Acknowledgments

We thank Fundação Oswaldo Cruz (Fiocruz) for provide part of the standard strains.

We thank Renata Helena Candido Pocente, Valdes Roberto Bollela, Gilberto Gambero Gaspar, and Roberto Martinez, from Hospital das Clínicas of Ribeirão Preto Medical School - University of São Paulo, for provide clinical isolates.

We thank Jussara Kasuko Palmeiro, from Universidade Federal de Santa Catarina, and Libera Maria Dalla-Costa, from Faculdades Pequeno Príncipe, Instituto de Pesquisas Pequeno Príncipe, for provide clinical isolates.

We thank Marcelo Dias Baruffi, from School of Pharmaceutical Sciences of Ribeirão Preto (FCFRP) – University of São Paulo (USP), Department of Clinical Analyses, Toxicology and Food Science (DACTB), for provide standard strains.

We thank Maria Regina Torqueti Toloi, Marcia Regina von Zeska Kress, Marcelo Dias Baruffi, and Andréia Machado Leopoldino, from FCFRP-USP, DACTB, for laboratorial support.

We also thank National Council for Scientific and Technological Development (CNPq, Brazil), Coordenação de Aperfeiçoamento de Pessoal de Nível Superior (CAPES, Brazil) and São Paulo Research Foundation (FAPESP, São Paulo State, Brazil) for all support in our research.

## Funding

Coordenação de Aperfeiçoamento de Pessoal de Nível Superior (CAPES, Brazil) Grant 8887.369851/2019-00 (NA). This study was financed in part by the CAPES - Finance Code 001. Pró-Reitoria de Pesquisa da USP Grant 18.1.796.60.2 grupo 057 (LNA).

Fundação de Amparo à Pesquisa do Estado de São Paulo (FAPESP, São Paulo State, Brazil), and by the Instituto Nacional de Ciência e Tecnologia Translational em Medicina (INCT-TM) grant CNPq/FAPESP 14/50891-1 (JECH).

University Global Partnership Network (UGPN) – Global Priorities in Cannabinoid Research Excellence Program (JAC).

National Council for Scientific and Technological Development (CNPq, Brazil) (JAC, JECH, and AWZ).

## Data and materials availability

All data associated with this study are available in the main text or the supplementary materials.

---

**Fig. S1.**
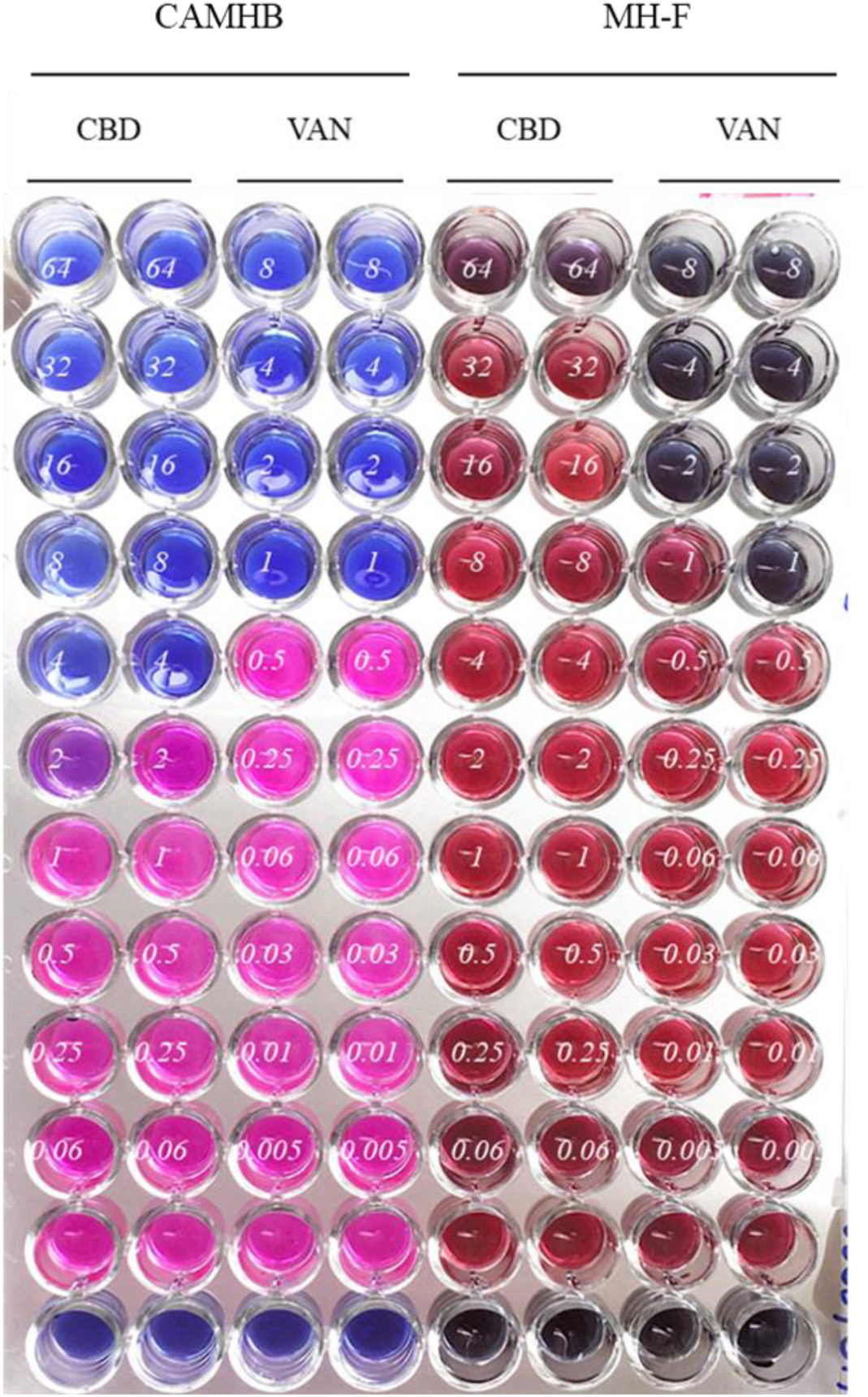
CBD MIC results against *S. aureus* ATCC 29213 observed from broth microdilution assay using MH-F broth and using CAMHB. We observed increased CBD MIC (64 µg/mL) when the assay was performed using MH-F broth (5% lysed horse blood + 0.1% β-Nicotinamide adenine dinucleotide [β-NAD] 20 mg/mL), in comparison with standard protocol using CAMHB for *S. aureus* (CBD MIC = 4 µg/mL). Increased MIC was also observed for vancomycin (VAN) control.

**Fig. S2.**
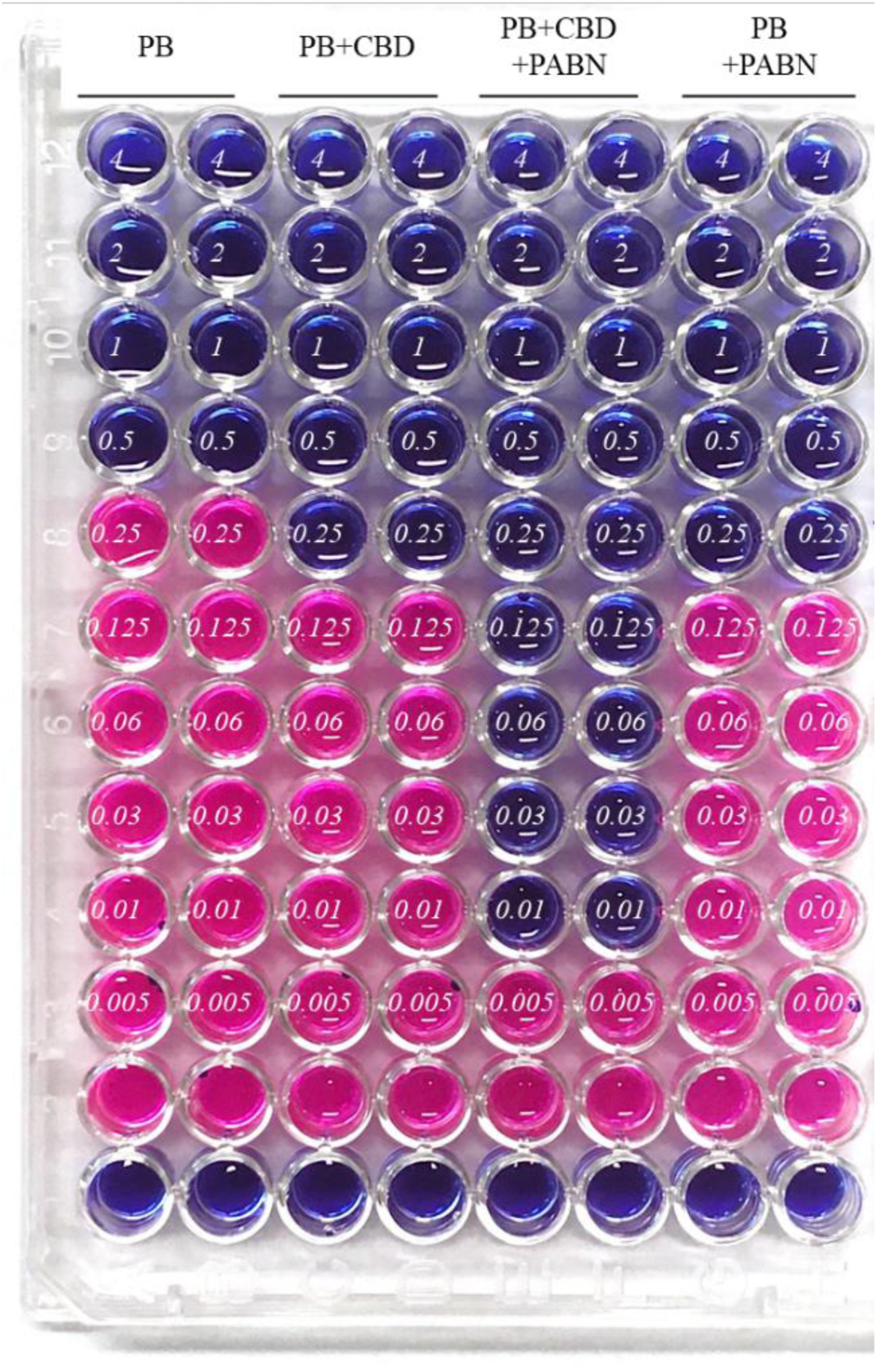
Representative screening assays results of antibacterial activity of the combination CBD + PB against PB-susceptible *K. pneumoniae* BAA-1705. The columns named “PB” are serial dilution of PB with no addition of CBD (PB MIC determination). The columns named “PB+CBD” are serial dilution of PB, plus a fixed concentration of CBD (256 µg/mL). The columns named “PB+CBD+PAβN” are serial dilution of PB, plus a fixed concentration of CBD (256 µg/mL) and a fixed concentration of PAβN (50 µg/mL). Finally, the columns named “PB+PAβN” are serial dilution of PB plus a fixed concentration of PAβN (50 µg/mL), with no addition of CBD. Blue wells show bacterial growth inhibition, while pink wells show bacterial growth. In each well, the numbers in italic refer to PB concentrations. Line “1 is MHB sterility control, while line “2” are bacterial growth control.

**Table S1.**
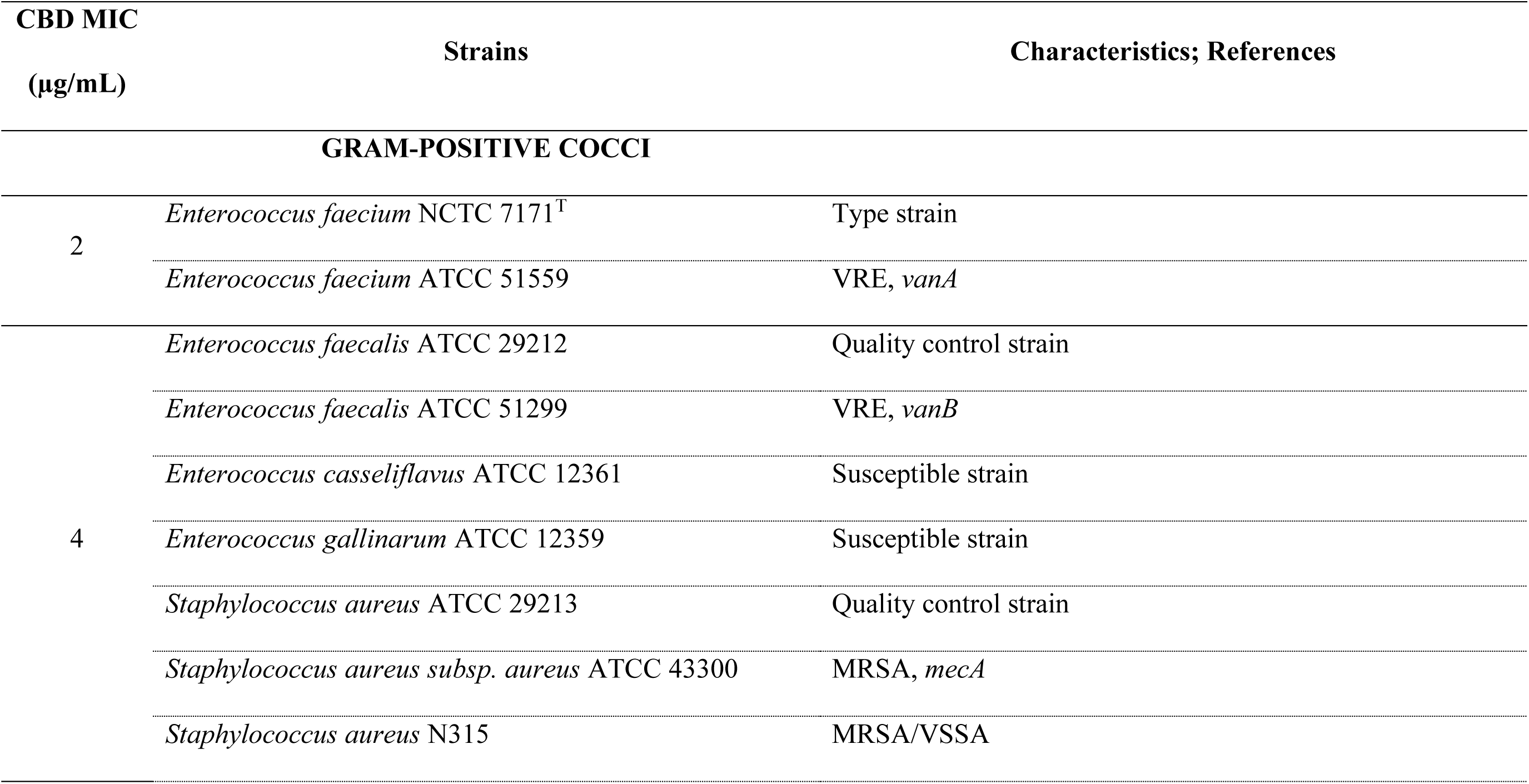
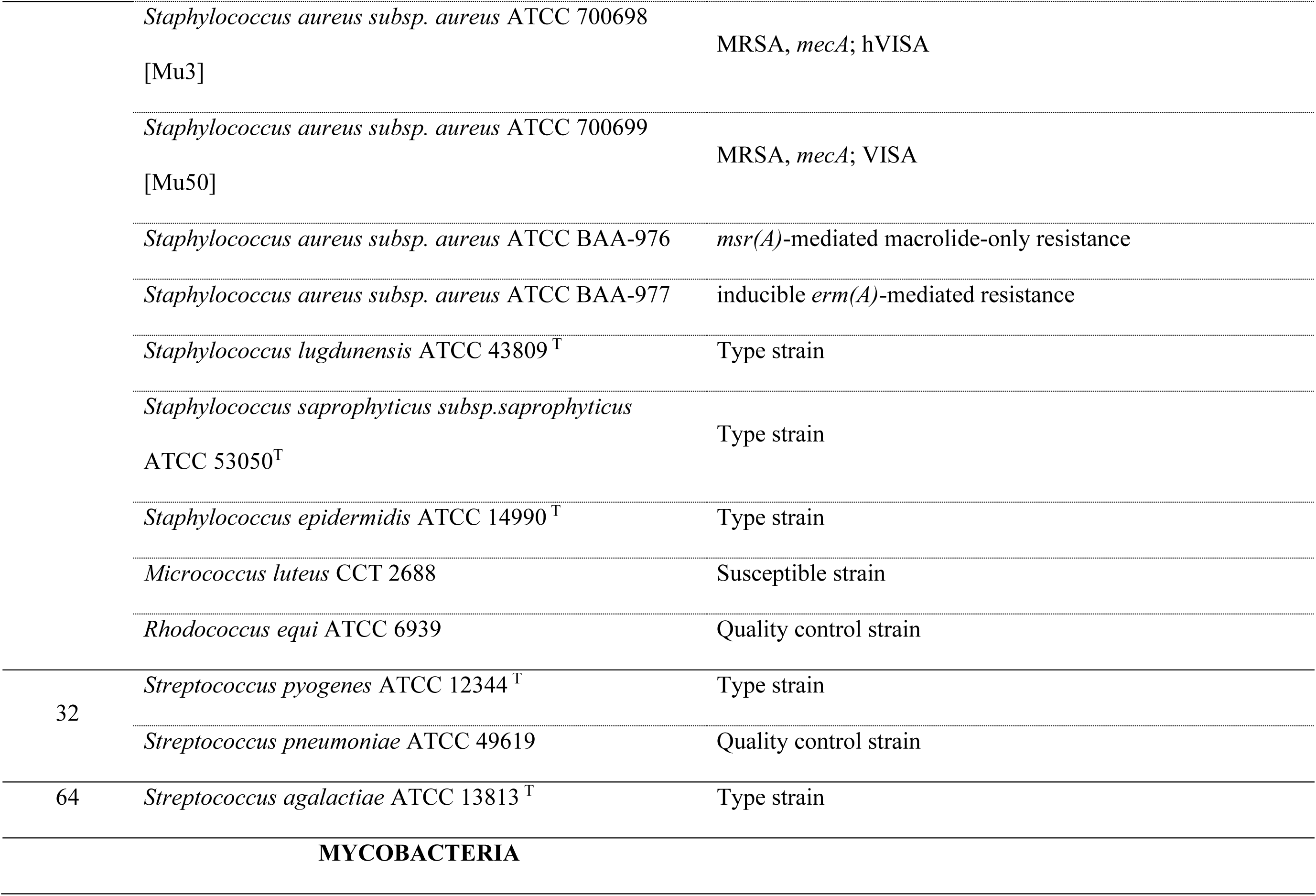
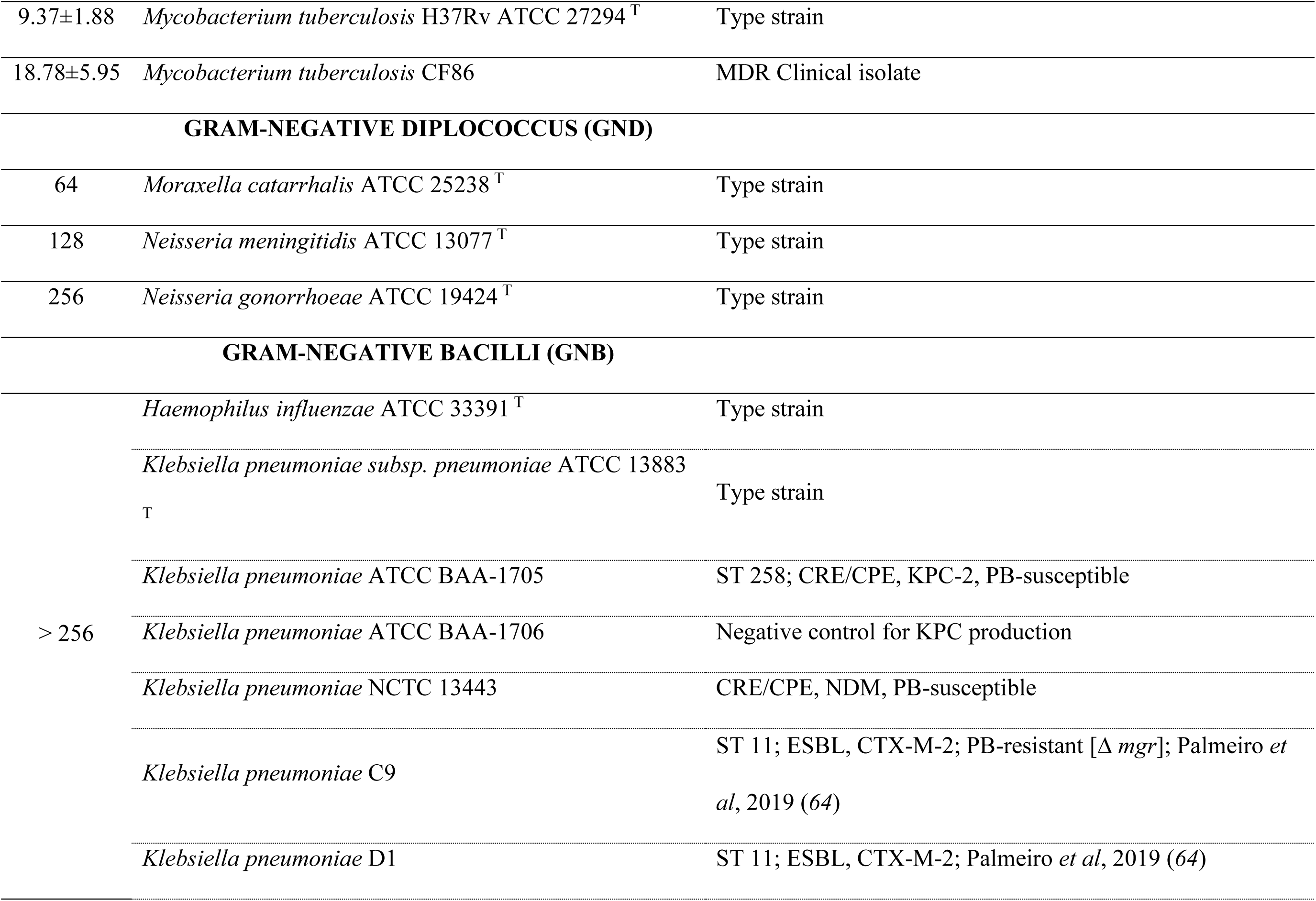
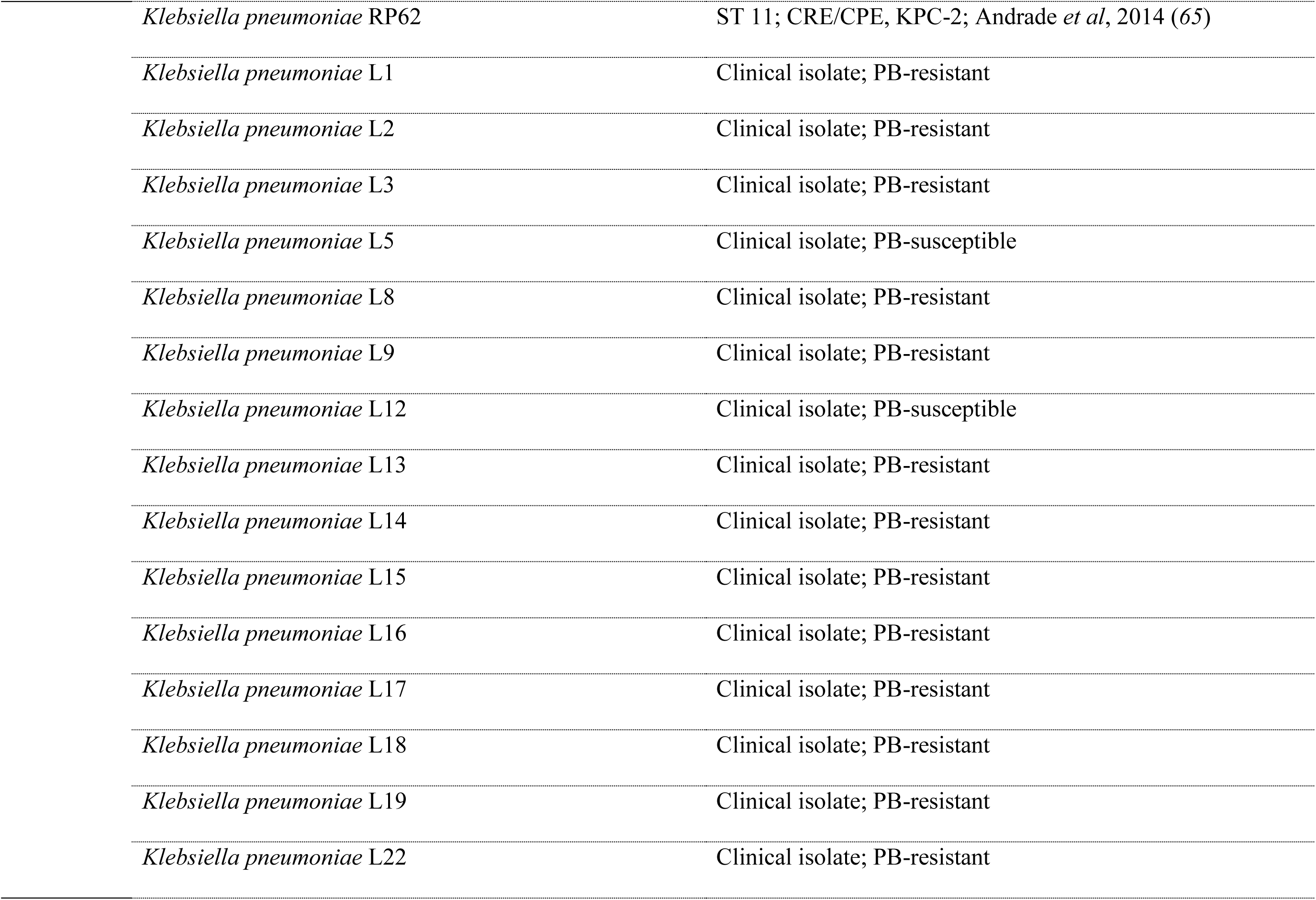
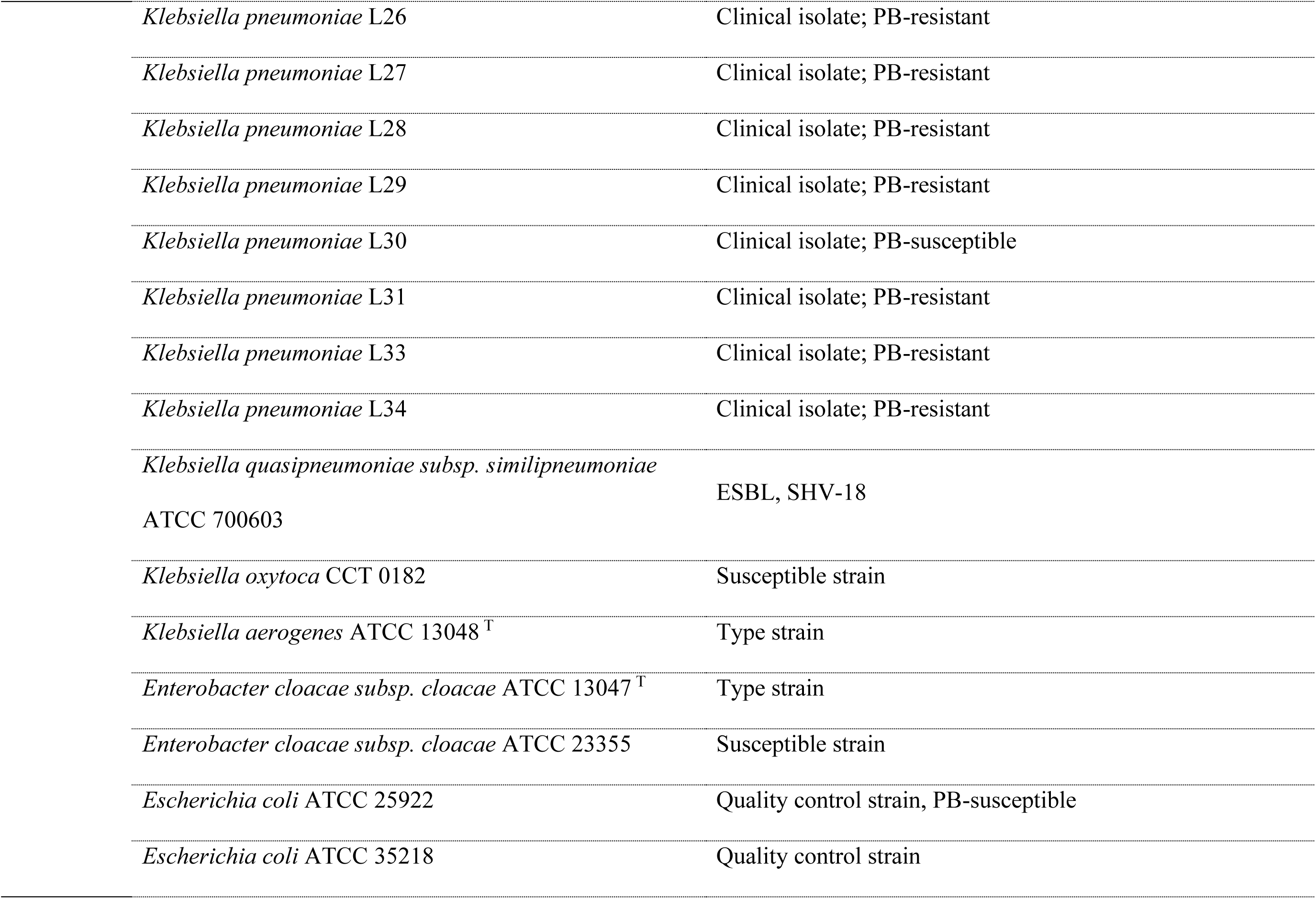
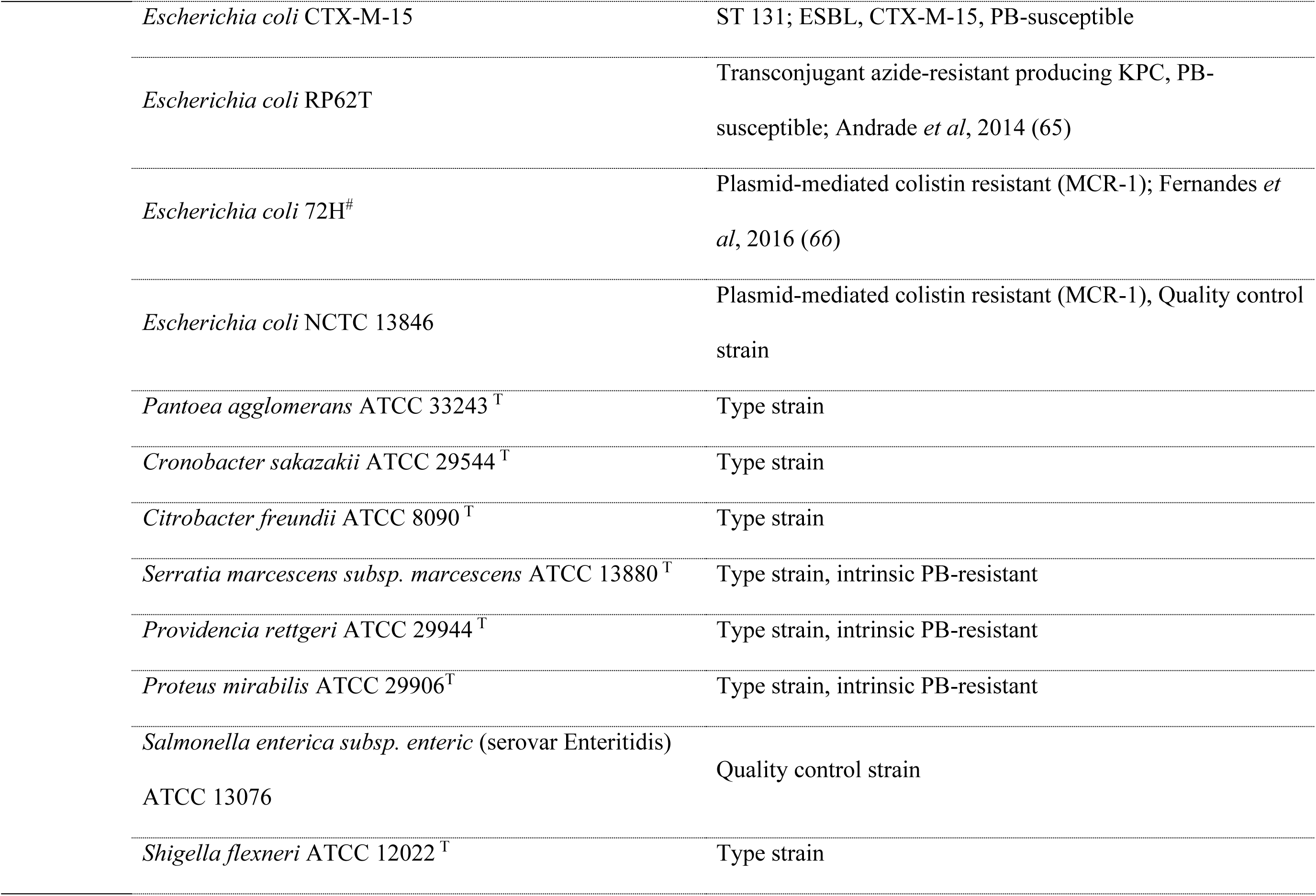
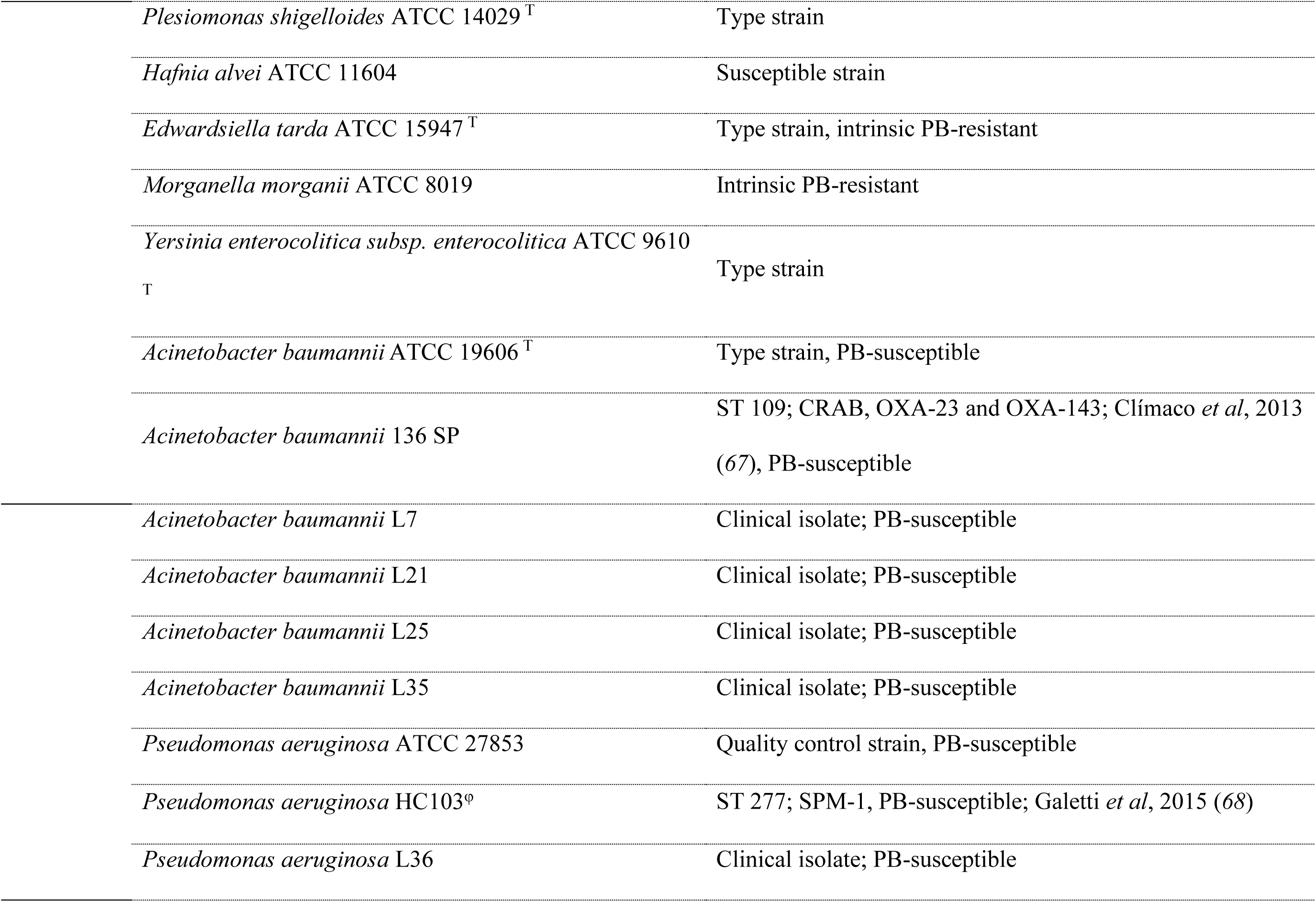
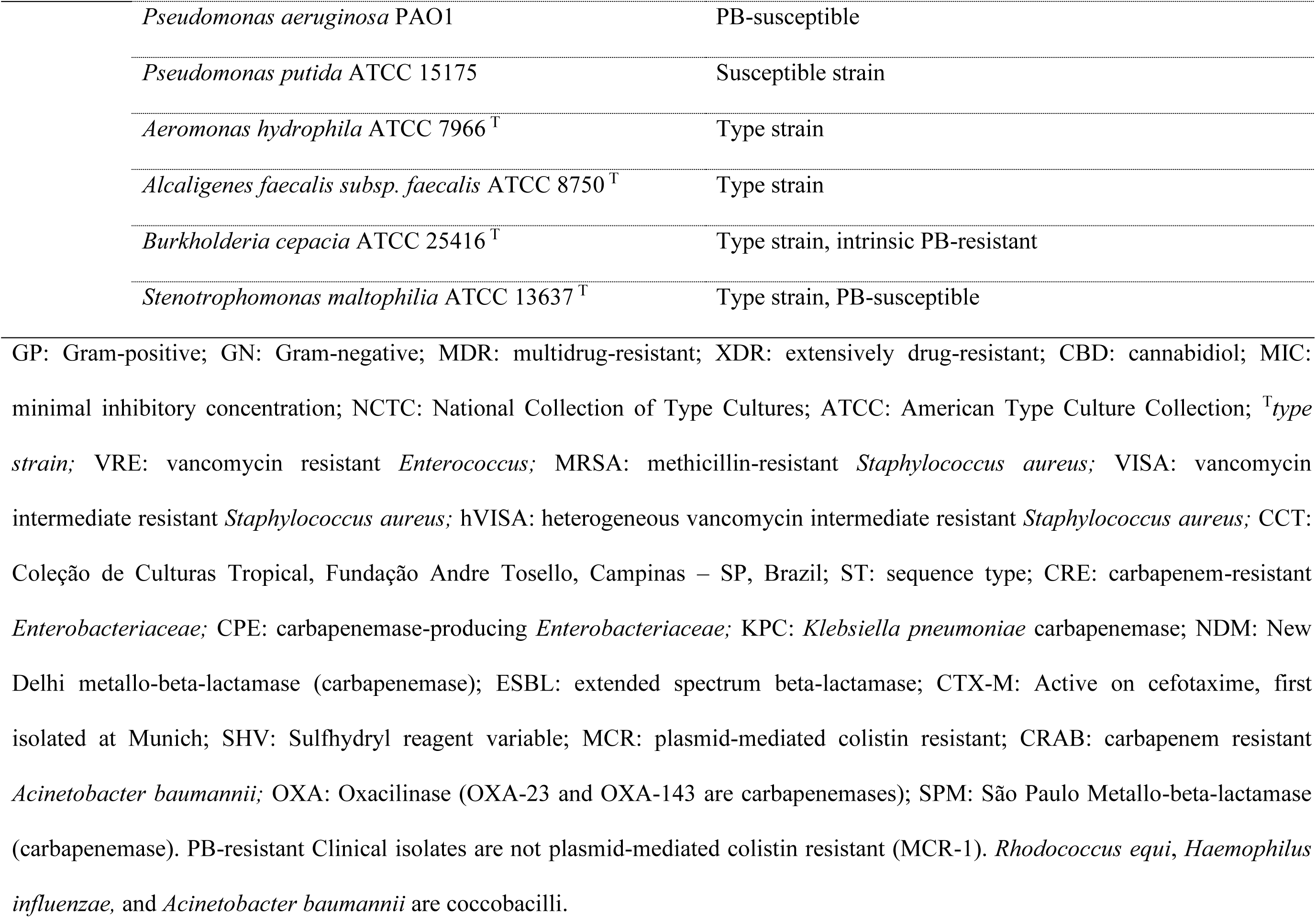
CBD MIC results against bacterial strains studied. Antibacterial activity of CBD was investigated against a broad panel of different bacterial species, comprehending GP (13 different species; 21 strains) and GN (30 different species; 73 strains) bacteria and *M. tuberculosis* (2 strains), including type-strains, quality control strains, and clinical isolates (including MDR strains, international high-risk clones, and susceptible strains).

